# VIRGO2: Unveiling the Functional and Ecological Complexity of the Vaginal Microbiome with an Enhanced Non-Redundant Gene Catalog

**DOI:** 10.1101/2025.03.04.641479

**Authors:** M. T. France, I. Chaudry, L. Rutt, M. Quain, B. Shirtliff, E. McComb, A. Maros, M. Alizadeh, F. A. Hussain, M. A. Elovitz, D. A. Relman, A. Rahman, R. M. Brotman, J. Price, M. Kassaro, J. B. Holm, B. Ma, J. Ravel

## Abstract

Despite the importance of the cervicovaginal microbiome, the mechanisms that govern its composition and drive its impact on host physiology remain poorly understood. This study expands our understanding of the function and ecology of the vaginal microbiome using VIRGO2, an enhanced non-redundant gene catalog comprising over 1.7 million well-annotated genes from body-site specific microbes and viruses. Analyses using VIRGO2 revealed novel insights, including the identification of previously uncharacterized vaginal bacteria, features of the vaginal mycobiome and phageome, and differential expression of bacterial carbohydrate catabolic genes. Constructed from over 2,500 metagenomes and 4,000 bacterial genomes, VIRGO2 broadens geographic representation and microbial diversity compared to its predecessor. This updated catalog enables more precise profiling of taxonomic and functional composition from metagenomic and metatranscriptomic datasets. VIRGO2 is a critical resource for integrative analyses of vaginal microbial communities and their interactions with host tissues, thereby enhancing our mechanistic understanding of vaginal health and disease.

## Introduction

Human microbiome science has progressed from identifying the resident microbes associated with health to characterizing how these microbes interact as communities within the host and contribute to host physiology. These microbial communities live in close association with host tissues and can affect physiologic change via direct interactions^1,2^ or through the production of metabolites and secreted products^3–5^. A mechanistic understanding of the interactions between host and microbes can enable the manipulation of this system to improve an individual’s health^6–9^. In pursuit of this mechanistic understanding, shotgun metagenomic and metatranscriptomic data are often used to describe the genomic content and *in vivo* transcriptional activity of the communities, respectively^10,11^. These techniques generate sequence data from bulk extracted DNA or RNA to provide an untargeted assessment of the genomes and transcriptomes that comprise microbial communities. However, the analysis and integration of these data is no trivial task. Sequences originating from tens to hundreds of microbial species might be found in each metagenome and metatranscriptome depending on the complexity of the underlying community. Reference-guided approaches that match sequence reads to genomic databases provide an attractive strategy to simplify the analysis of these complex data^12^. Yet, these approaches can yield unsatisfactory results when the reference database poorly matches the microbiome being analyzed^13^. Ecosystem-specific databases that are tailored to a particular habitat and microbial community can provide a more robust analysis of metagenomic and metatranscriptomic data^14^ and have been published for several different host site-specific microbiomes^15–19^. Here, we present VIRGO2, an enhanced iteration of the non-redundant gene catalog for the human vaginal microbiome, VIRGO (hereafter referred to as VIRGO1) ^20^, and demonstrate its use in gaining functional and ecological insight into this important microbiome with the re-analysis of 2,560 metagenomes and 363 metatranscriptomes.

The microbial communities that inhabit the human vagina are distinct from those found at other body sites, where community diversity roughly correlates with health^21^. Instead, an optimal vaginal microbiota (VMB) is often characterized by dominance of a single species of *Lactobacillus* and is associated with decreased risk for several adverse health outcomes (reviewed in France, et al. ^22^) including the acquisition of sexually transmitted infections^23–25^, spontaneous preterm birth^26,27^, and pelvic-inflammatory disease^28,29^. Lactobacilli are thought to dampen inflammatory responses^24,30,31^ and to exclude vaginal pathogens by lowering vaginal pH through their production of lactic acid^32^. A non-optimal VMB, on the other hand, is comprised of a more diverse and even mixture of facultative and obligate anaerobic bacterial species in the genera *Gardnerella*, *Prevotella*, *Fannyhessea*, *Sneathia*, *Ca.* Lachnocurva vaginae and *Megasphaera*, among others^33^. Women with these communities are often diagnosed with bacterial vaginosis, a common vaginal condition clinically defined by an elevated pH with a concomitant thin and homogenous discharge, fishy odor, and the presence of clue cells^34^. Yet, many women with a non-optimal VMB do not report experiencing any of these signs and symptoms^35^, but still appear to be at risk^36–38^. The composition of the VMB can also change over time, including shifting from an optimal *Lactobacillus*-dominated state to a non-optimal state, or the reverse^39–41^. Understanding the factors that govern the composition and functions of the VMB, and the mechanisms by which vaginal microbes influence host tissues, will enable the development of novel therapeutics and treatment options to promote vaginal health.

The VMB can be subdivided by taxonomic composition into seven community state types (CSTs)^42^, four of which are dominated by lactobacilli: I-*L. crispatus*, II-*L. gasseri*/*L. paragasseri*, III-*L. iners*, V-*L. jensenii*/*L. mulieris*. The remaining three CSTs (IV-A, IV-B, IV-C) are characterized by a diverse mixture of facultative and obligate anaerobes; IV-A is distinguished by the presence of *Ca*. L. vaginae (formerly BVAB1), while IV-B is characterized by higher relative abundance of *Gardnerella* spp. and IV-C by the absence of *Gardnerella* and *Fannyhessea*. Of the four *Lactobacillus*-dominant CSTs, only CST I, II, and V are considered optimal. *L. iners*-dominated communities shift to CST IV at greater rates than those dominated by other lactobacilli, and do not appear to offer the same degree of protective benefits to the host^39,43–46^. Recent studies of the VMB have utilized shotgun metagenomics and metatranscriptomics to describe the vaginal microbiome and have revealed additional complexity not captured by the CST paradigm. For example, in the release of VIRGO1, we highlighted that vaginal bacterial populations appear to not be genetically homogenous, and instead comprised of multiple genotypes^20^. Since then, we have leveraged this intraspecies diversity to identify more complex microbial population configurations that are common across women and are associated with symptomatic BV^47^. Other studies have found that genetic diversity within *Gardnerella* is associated with preterm birth^27,48^. Analyses of the vaginal metatranscriptome revealed further complexity by demonstrating that the transcriptional activity of a bacterium is not always correlated with its abundance in a community^49,50^. Still more studies have integrated metagenomic data with metabolomic data to identify and characterize enzymatic differences between *L. iners* and other vaginal lactobacilli^51,52^. From all these studies, it has become clear that integrative multi-omic analyses are needed to fully understand the vaginal microbiome and its relationship to health.

Here, we describe the construction, validation and application of VIRGO2, an enhanced iteration of a non-redundant gene (NRG) catalog for the vaginal microbiome. Since its release, VIRGO1 has seen regular use in the analysis of vaginal metagenomic and metatranscriptomic data and has proven to be a powerful tool for analyzing and integrating these data types. VIRGO2 was constructed *ab initio* from a large sequence dataset of >2,500 shotgun metagenomes and >4,000 whole genome sequences from relevant bacteria, fungi, viruses, and protists. For this version, we emphasized broadening the geographic representation of the hosts from which sequence data were collected to build the catalog and improving VIRGO’s ability to characterize the non-bacterial components of the vaginal microbiome. The resulting NRG catalog is approximately twice the size of the original and was validated for its ability to characterize the vaginal microbiome. A re-analysis of vaginal metagenomes and metatranscriptomes demonstrated how VIRGO2 can be used to gain insight into the VMB and highlighted novel ecological and functional findings related to the bacteria, bacteriophages, and fungi that inhabit the human vagina.

## Results

### *De novo* assembly and binning of shotgun metagenomes to generate MAGs

The majority of the sequence data used to construct VIRGO2 was sourced from 2,560 shotgun metagenomes derived from vaginal (n=2,496), or penile urethral (n=64) samples (see Supplementary Table 1 for accession numbers; metagenomes originate from previously published studies^23,50,53–62^). The small collection of penile metagenomes was derived from a North American sexual network study^55^ and was included to more comprehensively characterize the organisms that might be identified in vaginal ‘omics data. In total, the metagenomes were derived from samples provided by 2,048 individuals located in six different countries and representing five different continents (Fig. 1a). All samples were provided by reproductive-age individuals, and 1,173 (47%) of the vaginal metagenomes were derived from samples provided by pregnant women. The metagenomes from North America derived from pregnant and non-pregnant women. All other continents were represented by samples from either only non-pregnant women (Fiji) or only pregnant women (Europe, Asia, Africa).

**Fig. 1.**
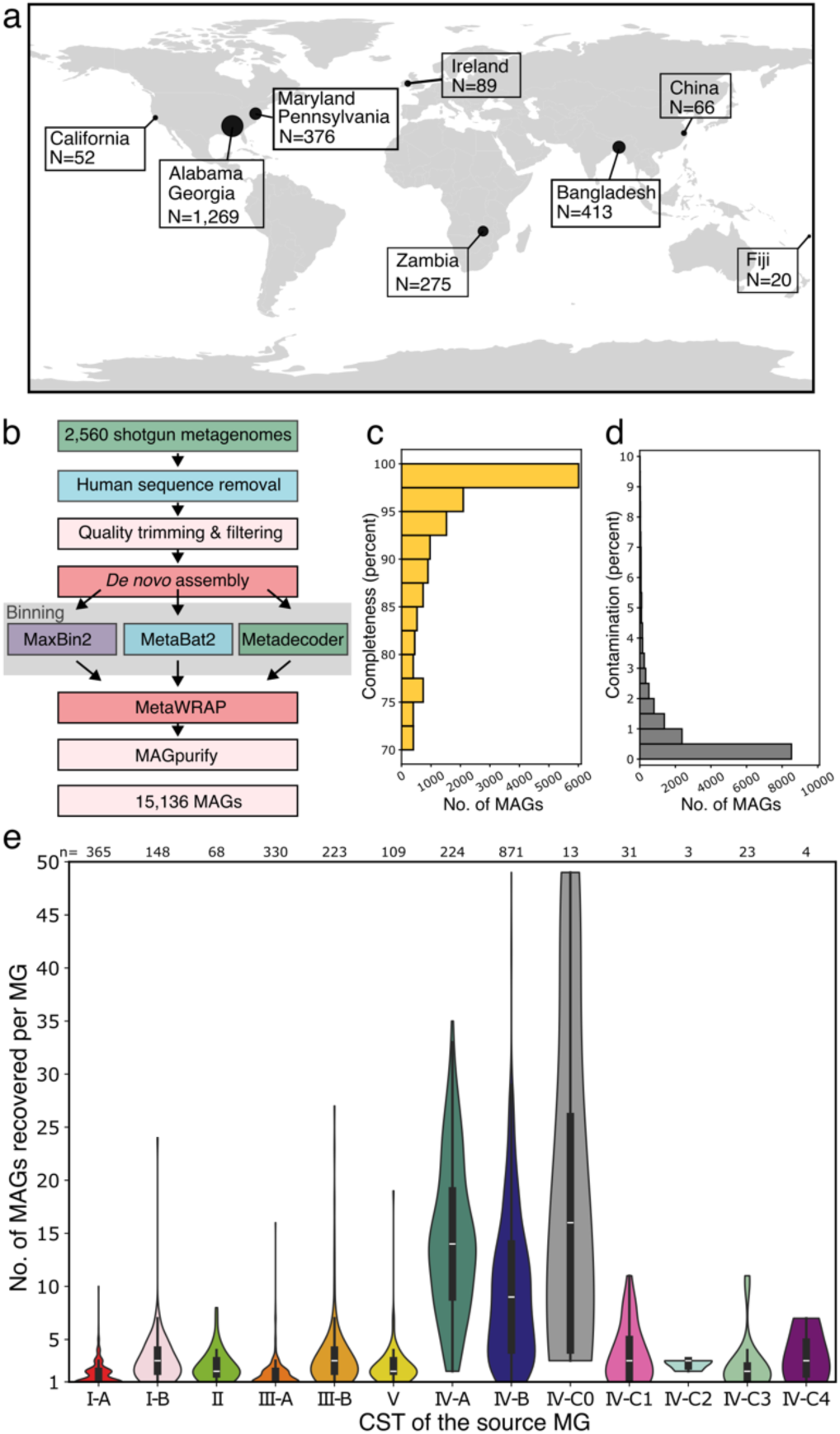
Overview of the metagenomes, metagenome-assembled genomes, and bioinformatics processes used to construct VIRGO2. **a,** Geographic origin of shotgun metagenomes used in the construction of VIRGO2 (see Supplementary Table 1 for accession numbers). **b,** Bioinformatics pipeline used to generate metagenome assembled genomes from the shotgun metagenomes. **c,** CheckM completeness and **d,** contamination estimates of the 15,136 MAGs generated in this study. **e,** Association between the number of MAGs recovered and the community state type of the metagenome. Numbers above the violin plots denote the number of metagenomes assigned to each CST.

Each metagenome was processed using a *de novo* assembly and binning pipeline (Fig. 1b) to generate metagenome assembled genomes (MAGs). This pipeline produced a total of 10,467 high and 4,670 medium quality MAGs, of which 10,598 (70.0%) were estimated to be more than 90% complete (Fig. 1c) and 14,591 (96.4%) were estimated to be less than 5% contaminated (Fig. 1d). Many of medium quality MAGs that were estimated to have lower completeness represented poorly characterized species (Supplementary Table 2, e.g. *Ca*. L. vaginae, average completeness 75.8%). Almost all the metagenomes produced at least one MAG (n=2,412; 94.2%), and many produced several (range: 0-48, median: 3). Not surprisingly, the number of MAGs produced from a metagenome was found to vary with community state type. Metagenomes that were dominated by single species yielded fewer MAGs than those that were more compositionally diverse and even (Fig. 1e; e.g. CST I-A vs IV-B, Poisson regression, p_adj_ < 0.01).

### A comprehensive inventory of the bacteria that inhabit the human vagina

The collection of 15,137 MAGs represents a broad and unbiased sampling of the bacterial species which inhabit the vaginal microbiome. The taxonomy of the MAG collection was established using the Genome Taxonomy Database Toolkit (GTDB-tk), revealing 288 unique bacterial species, including 146 that were present in at least five metagenomes. Among these, *Lactobacillus iners* was the most prevalent species identified and was found in 1,380 metagenomes (53.9%, Fig. 2a). *L. iners* MAGs were routinely produced from communities where it was the dominant species, but also from communities where it was co-resident with other lactobacilli or with non-*Lactobacillus* species (Fig. 2a, radar plot). Other familiar prevalent species include *F. vaginae* (formerly *Atopobium vaginae*, 27.6%), *M. lornae* (25.9%), *L. crispatus* (24.1%), *Amygdalobacter indicium* (formerly BVAB2^63^, 19.6%), *Ca.* L. vaginae (formerly BVAB1, 15.1%), as well as several species in the genera *Gardnerella*, *Prevotella*, and *Fannyhessea* (Fig. 2a). Our analysis also revealed some prevalent species that, although recognized in the GTDB, have yet to be given names, including: *Berryella* sp001552935 (21.8%), *Dialister* sp001553355 (15.9%), *Parvimonas* sp001552895 (11.9%), and *Prevotella sp000758925* (9.3%). All four of these bacteria were essentially only identified in communities that were not dominated by *Lactobacillus* (Fig. 2a, radar plots).

**Fig. 2.**
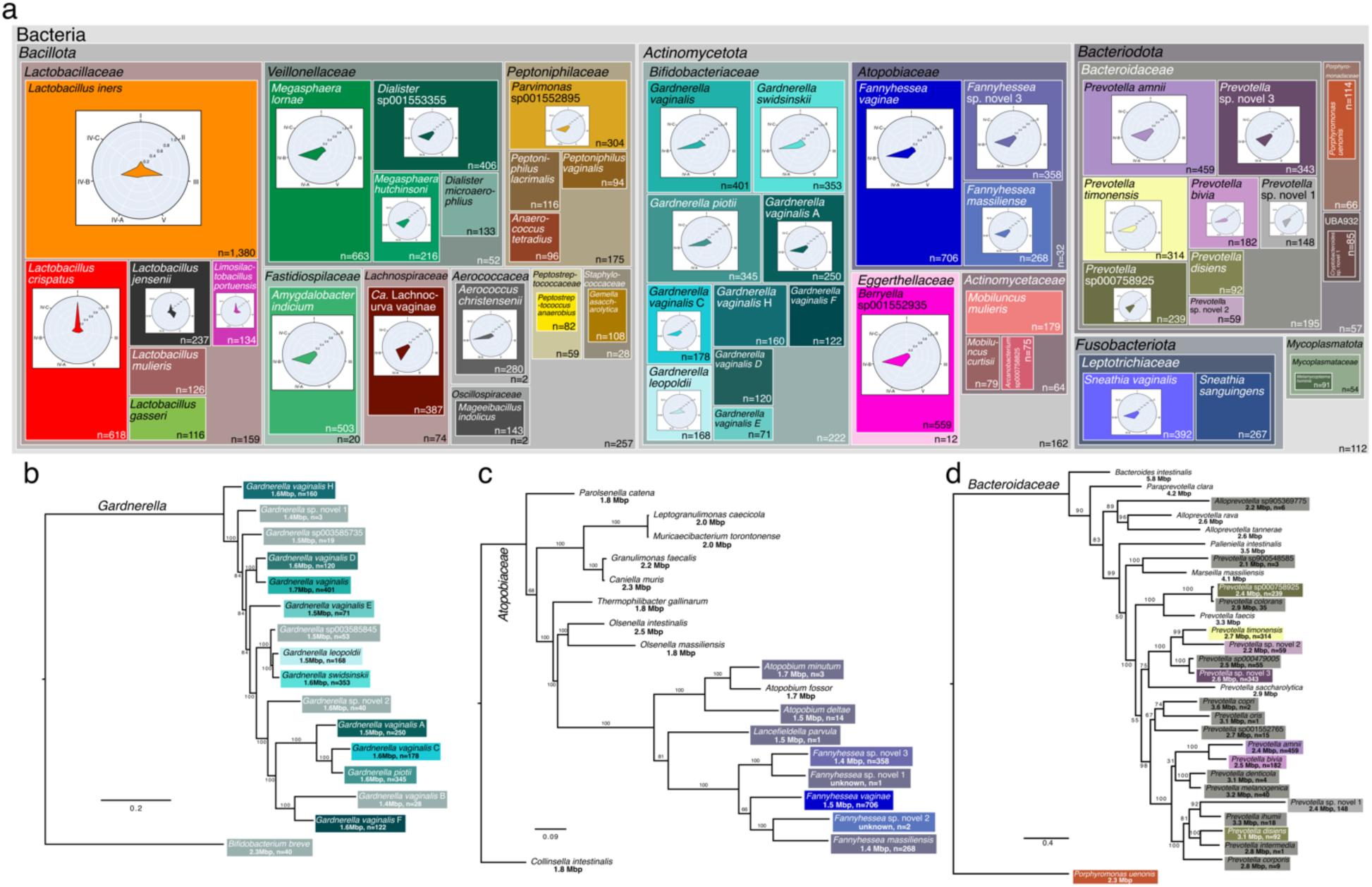
Description of the taxonomy and prevalence of vaginal bacteria. **a,** Treemap diagram displaying the number of MAGs generated for each of the 50 most prevalent species. Empty spaces within phyla represent the number of residual MAGs in that phylum not accounted for by the displayed species. Interior radar plots indicate the CST distribution of the metagenomes that originated the species’ MAGs. Concatenated core genome phylogenetic trees of the genera where prominent novel species were identified including: **b,** *Gardnerella,* **c,** *Fannyhessea* and **d,** *Prevotella*. Leaves are labeled with the species’ designation along with the species’ average genome size and the number of MAGs generated.

The collection of MAGs also included many genomes which could not be assigned to a species by the GTDB-tk, indicating they may belong to novel or poorly characterized species. To classify these genomes, we first collected representative genomes of closely related species and then performed a phylogenetic analysis. Two novel species were identified in the genus *Gardnerella*, increasing the total number of *Gardnerella* species to 15^64^. Neither novel *Gardnerella* species were particularly prevalent, being identified in just 3 (0.1%) and 40 (1.6%) metagenomes respectively (Fig. 2b). We have since isolated the second of these novel *Gardnerella* species from a vaginal swab specimen and confirmed its identity via whole genome sequencing (ascn: ACNEUN). Three novel species were identified in *Fannyhessea* (Fig. 2c), two that were rare (*Fannyhessea* sp. novel 1, n=1; *Fannyhessea* sp. novel 2, n=2), and one that was common (*Fannyhessea* sp. novel 3, n=358, 14.0%). We found *Fannyhessea.* sp. novel 3 to share a clade with *Fannyhessea* sp. novel 1 whereas, *Fannyhessea* sp. novel 2 was more closely related to *F. vaginae* and *F. massiliensis* (Fig. 2c). Another three novel species were identified in *Prevotella*, all of which were fairly common. *Prevotella* sp. novel 1 was identified in 148 metagenomes (5.8%) and is related to *P. ihumii* (Fig. 2d). The remaining two novel *Prevotella* species share a clade with *P. timonensis* and *P.* sp000479005, and of these four, *Prevotella* sp. novel 3 was the most frequently identified (n=343, 13.3%), surpassing even *P. timonensis* (n=314, 12.2%).

### Construction and annotation of the VIRGO2 non-redundant gene catalog

To construct the VIRGO2 gene catalog, the 15,136 MAGs were supplemented with sequence data from: 1) the residual un-binned metagenome contigs (length >2,000bp) and 2) isolate whole genome sequences (WGS) from relevant bacteria, viruses, fungi, and protists (254 unique species, 4,013 genomes, accession numbers in Supplementary Table 3). Prodigal identified 21,202,535 coding sequences (CDS) from the MAGs, 8,294,162 from the residual un-binned contigs and 8,015,873 from the isolate WGS (total=37,512,570 CDS). The CDS were then clustered using CDhit (95% sequence identity and 90% overlap), and for each cluster, the longest gene was selected as a representative, producing 1,773,155 non-redundant genes (NRGs), roughly a doubling of that included in the VIRGO1 gene catalog (∼1.8M versus ∼0.9M). Around one third of the NRGs were unique (cluster size of 1, n=673,407) and the remaining two thirds had a median cluster size of 5 (Supplementary Fig. 1). The NRGs were next annotated with taxonomic and functional information to facilitate their use in characterizing vaginal microbiome ‘omics data. The taxonomic annotation of each NRG was derived from that of the source sequences in the corresponding gene cluster (Fig. 3a). In total, taxonomic annotations were applied to 91% of the NRGs in VIRGO2, which is a dramatic improvement over VIRGO1 (47%). Functional annotations were also applied to the NRG using eggNOG-mapper^65^ (v2, default settings), PANNZER2^66^, and AMRFinderPlus^67^. The applied functional schemes included eggNOG^68^, COG category^69^, PFAM^70^, GO terms^71^, Enzyme Commission numbers^72^, BiGG Reaction^73^, KEGG^74^, and Carbohydrate Active Enzymes^75^, Gene Product^66^, antimicrobial resistance genes^67^, and secondary metabolite biosynthetic gene pathways^76^. Combined, 81% of the VIRGO2 NRGs have at least one functional annotation (Supplementary Fig. 2).

**Fig. 3.**
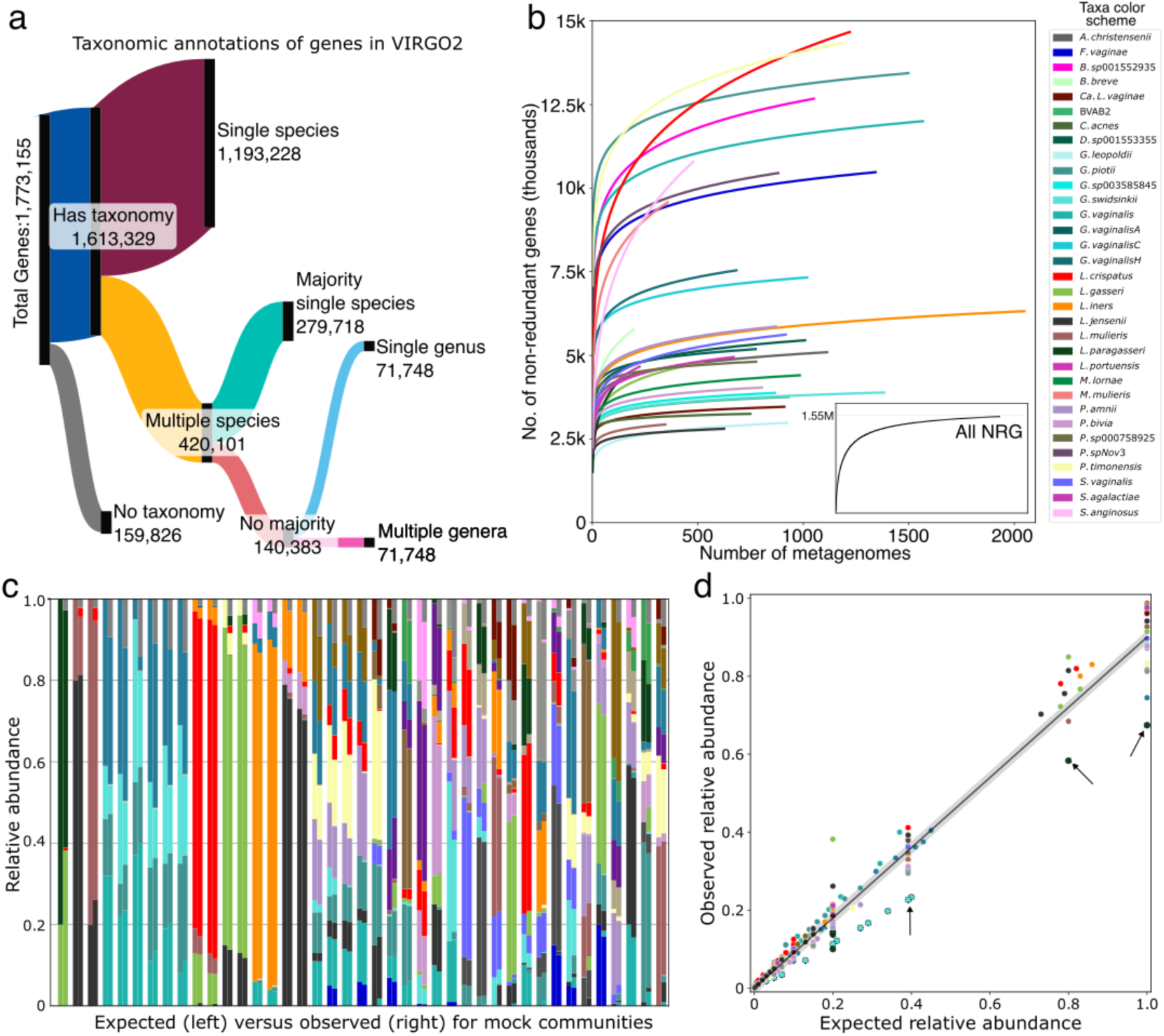
Annotation and validation of the VIRGO2 non-redundant gene catalog. **a,** Sankey diagram displaying the taxonomic assignment decision tree for VIRGO2 non-redundant genes. **b,** Gene accumulation curves showing the change in the number of non-redundant genes for a species as the number of metagenomes containing that species increases. Inset graph shows the accumulation curve for the overall number of non-redundant genes in VIRGO2. **c,** Comparison of the expected (left) and VIRGO2 estimated (right) taxonomic compositions of mock shotgun metagenomes. **d,** Relationship between expected and observed relative abundances (linear regression, r^2^=.988, p<0.001). Arrows indicate instances where VIRGO2 is slightly underestimating the relative abundances of *G. swidsinskii* and *L. paragasseri*.

### Validation of VIRGO2 demonstrates comprehensiveness and accuracy

VIRGO2 is intended to comprise a union of pangenomes of the microbes that inhabit the vagina, i.e., the vaginal microbiome’s “meta-pangenome”^77^. To test if VIRGO2 adequately encompasses each species’ genetic diversity, we generated NRG accumulation curves that track the total number of NRGs with increasing numbers of metagenomes. For the catalog overall (Fig. 3b, inlay), and for most of the individual species we examined, the curves demonstrated a rapid saturation in the repertoire of NRGs (Fig. 3b). However, for some species the accumulation curves failed to approach an asymptote, indicating that VIRGO2 may benefit from the inclusion of data from more samples containing these species. Among those not reaching saturation were *L. crispatus* and *P. timonensis*, two prominent vaginal bacteria that were well-represented in the metagenomes used to build VIRGO2. To define the scope of genetic diversity in these two species, we grouped their NRGs into orthologous gene clusters and generated pangenome accumulation curves. For both species, the accumulation curves approach saturation at ∼7000 orthologs (Supplementary Fig. 3), indicating that much of their diversity in NRG content originates from genetic variants of orthologous genes. These results demonstrate that VIRGO2 comprises the genetic and functional diversity of vaginal microbes.

We next sought to validate VIRGO2’s ability to characterize the taxonomic composition of vaginal microbiome ‘omics data. To do this, we constructed mock communities containing known mixtures of species using synthetic metagenomic sequence reads. Some of the mock communities were designed to mimic natural community structures, while others comprised random assortments, or specific combinations of closely related species (e.g. *L. jensenii* and *L. mulieris*, Fig. 3c). The synthetic reads were mapped to the VIRGO2 NRG catalog and taxonomy was estimated as relative abundances (corrected for gene length). We found that VIRGO2 accurately characterized community composition even when presented with particularly challenging cases (Fig. 3c). For example, VIRGO2 was able to disentangle the relative abundances of closely related *Gardnerella* species when presented with simulated mixtures of varying proportions. One exception was that VIRGO2 appears to slightly underestimate the relative abundances of *G. swidsinskii* and *L. paragasseri* (Fig. 3d, arrows). Further investigation revealed that both species have close relatives (*G. leopoldii* and *L. gasseri*, respectively) that share some NRG clusters, causing them to be labeled at the genus, rather than species level. Nevertheless, the overall correlation between expected and observed relative abundances for these synthetic mock datasets was high (r^2^ 0.988, p<0.001, linear regression, Fig. 3d), indicating that VIRGO2 accurately characterizes vaginal microbial community composition.

### Taxonomic composition of the vaginal microbiota as characterized by VIRGO2

One of the main functions of VIRGO2 is to estimate the composition of vaginal microbiome ‘omics datasets. To demonstrate this, we characterized the taxonomic composition of the 2,560 metagenomes used in the construction of the catalog. Sequence reads were mapped to VIRGO2, and relative abundances were estimated as described for the validation data. Observed community compositions resembled those identified by prior 16S rRNA gene amplicon sequencing surveys but with substantial added complexity due to the enhanced taxonomic resolution provided by VIRGO2 (Fig. 4a). Our inclusion of metagenomes from multiple locations around the world allowed us to evaluate if vaginal bacteria display biases in their prevalence across geographies. We found that most bacterial species were common in metagenomes from every sampled continent, indicating they have cosmopolitan distributions (Fig. 4a). This includes both *L. crispatus* and *L. iners*, as well as most species that frequent CST IV communities (Fig. 4a). However, some species do exhibit biases in their geographic distribution. The sister species *L. gasseri* and *L. paragasseri* were both more prevalent in metagenomes from North American and European women than other geographies (Fig. 4a, Supplementary Table 4). Whereas *L. portuensis* was common in *L. crispatus* and *L. iners* dominated metagenomes from women in Asia but rare elsewhere (Fig. 4a, Supplementary Table 4). Finally, *Ca*. L. vaginae was rare in CST IV communities from Asia but common to these communities from other continents (Fig. 4a, Supplementary Table 4). These results indicate that while most vaginal bacterial species exhibit cosmopolitan distributions, there are several species that do display biases in their geographic distributions.

**Fig. 4.**
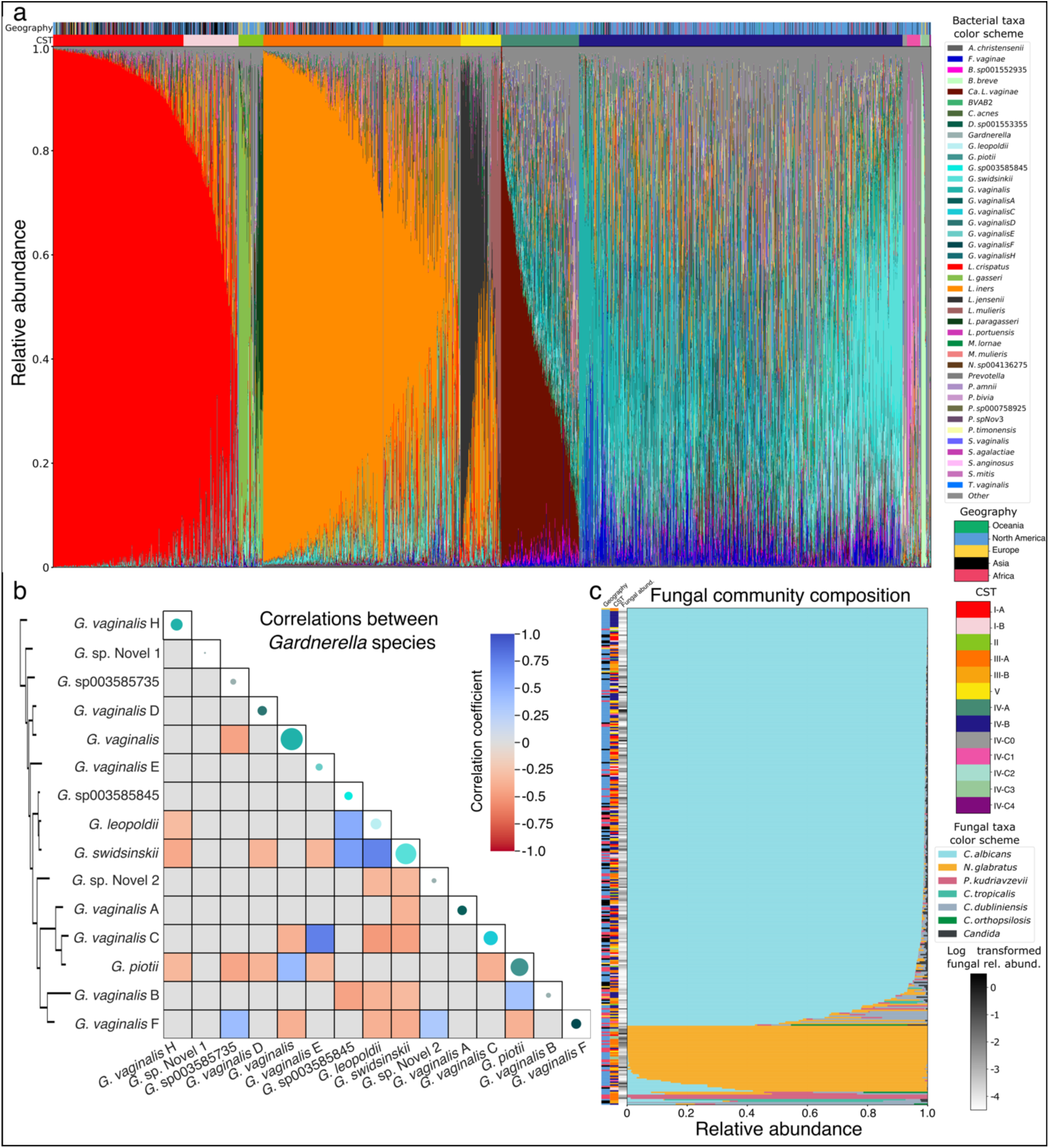
Description of the taxonomic composition of the vaginal microbiome and mycobiome, as estimated using VIRGO2. **a,** VIRGO2 estimated taxonomic composition of the 2,560 metagenomes used in its construction. The geographic origin and community state type of the metagenomes are displayed above the bars. **b,** Pairwise correlation coefficients in the observed relative abundances of *Gardnerella* spp. Coefficients were estimated using ANCOMBC, and only significant values are displayed (p<0.001). Colored points along the diagonal represent the average relative abundance of the species observed *in vivo*. The species are ordered by the phylogeny produced for Fig, 2b and that phylogeny is again displayed to the left of the panel. **c,** VIRGO2 estimated taxonomic composition of the mycobiome for the 339 (13.2%) metagenomes where fungi were detected. The sample’s geographic origin and the community state type of the corresponding microbiome are displayed to the left of the bars, along with a heatmap representing the log_10_ transformed total fungal relative abundance.

One major advancement provided by VIRGO2 is its ability to distinguish the now 15 recognized species of *Gardnerella*. Of the 15, *G. vaginalis*, *G. swidsinskii,* and *G. piotti* were the most common in the analyzed metagenomes (Fig. 4a). Unlike the vaginal lactobacilli, *Gardnerella* spp. are often identified together in the vaginal microbiome. The median number of *Gardnerella* species identified in a metagenome was four and the maximum was twelve. To determine if there was any structure to the subcommunities of *Gardnerella* spp., we estimated correlation coefficients between the relative abundances of the individual species. While most pairwise correlations between *Gardnerella* spp. were not significant, some significant positive correlations (n=8) and negative correlations (n=23) were identified (Figure 4b, p<0.001, ANCOMBC). Many of the identified correlations involved members of the subclade containing *G. swidsinskii*, *G. leopoldii*, and *G.* sp00358545. These three species had positive correlations with each other and negative correlations with many other *Gardnerella* species (Figure 4b) and *G. swidsinskii* alone accounted for eight of the observed negative correlations (34.7%. Subsequent investigation revealed that most *G. swidsinskii* genomes (88% of isolate WGS, 79% of MAGs) harbor a unique biosynthetic gene pathway for the production of a lacticin 481-like bacteriocin (Additional File 1). This bacteriocin could drive the numerous observed negative correlations between *G. swidsinskii* and other *Gardnerella* spp.

### The vaginal mycobiome as characterized by VIRGO2

A key new feature of VIRGO2 is its ability to characterize the vaginal mycobiome. During the construction of the catalog, genomic data of several fungal species were identified in the un-binned contigs of the metagenomes. These data were further supplemented with isolate whole genome sequences of relevant species. A total of 27,545 fungal NRGs are included in VIRGO2 belonging to six species including: *Candida albicans*, *Nakaseomyces glabratus* (formerly *C. glabrata*), *Pichia kudriavzevii* (formerly *C. krusei*), *C. tropicalis*, *C. dubliniensis* and *C. orthopsilosis*. To characterize the vaginal mycobiome, we extracted the relative abundances of these fungal species from the VIRGO2 mapping results. Fungi uncommonly comprise a significant proportion of the overall microbial community—only 339 (13.2%) of the metagenomes had a total fungal relative abundance over 10^-4^ (Fig. 4c). *L. iners*-dominated bacterial communities were over-represented (n=110, 38.6%) in the subset of fungal-containing metagenomes (Fig. 4c, logistic regression, p_adj_<0.01). We next examined the taxonomic composition of the mycobiome. *C. albicans* was the dominant fungal species in most of these metagenomes (n=285, 84.1%, Fig. 4c). A smaller subset of metagenomes had a mycobiome dominated by either *N. glabratus* (n=47, 13.9%) or *P. kudriavzevii* (n=3, 0.9%). Mycobiomes that were dominated by *N. glabratus* tended to represent a higher proportion of their overall metagenome than those that were dominated by *C. albicans* (Fig. 4c, anova, p_adj_=0.0045). However, both *C. albicans* and *N. glabratus* were found to occasionally represent a more substantial proportion of the overall metagenome (max 29.2% and 21.2%, respectively).

### Identification and characterization of bacteriophage in metagenomic data using VIRGO2

Bacteriophages, both as active virions and integrated prophages, are thought to be a significant component of the microbiome, but their abundance and role in vaginal ecology remains poorly described. To enable VIRGO2’s ability to characterize these viruses, we identified and annotated bacteriophage sequences in the metagenomic data used in the construction of the database. A total of 3,414 bacteriophage contigs encoding 89,793 of the NRGs were identified and then annotated with a taxonomy, lifestyle (temperate versus virulent), and predicted host (Supplementary Table 5). The relative abundance of bacteriophages was estimated using the VIRGO2 mapping results for the 2,560 metagenomes. We found bacteriophages typically represent 5-10% of the vaginal metagenome, with *L. crispatus* dominated communities having a slightly higher proportion of bacteriophages than other CSTs (Fig. 5a). However, there were a few instances where the proportion of bacteriophages was much higher (max observed 43.7%). The lifestyle annotations were then used to identify the 63 metagenomes (2.4%) where the proportion of virulent bacteriophages was greater than that of temperate (Fig. 5b). Four of these metagenomes originated from the same person, spread evenly over a 10-week longitudinal sampling period. The peak bacteriophage abundance occurred at Timepoint 4, when the bacterial community experienced a concurrent decline in the relative abundance of *L. gasseri* (Fig. 5c). We were able to visualize putative parts of a once intact bacteriophage by transmission electron microscopy (TEM) of filtrate of stored samples collected the day prior to Timepoint 4 (Fig. 5d). The ∼50kbp genome of the predicted virulent bacteriophage driving these observations was also identified in the metagenome assembly from the four timepoints and annotated using Pharokka (Fig. 5e). Lysogeny related genes were not identified in the genome, consistent with the predicted virulent lifestyle of the bacteriophage, and several genes encoding tail-fiber proteins were identified, consistent with the TEM micrograph. These results demonstrate how VIRGO2 can be used to identify and characterize bacteriophage populations in the vaginal microbiome.

**Fig. 5.**
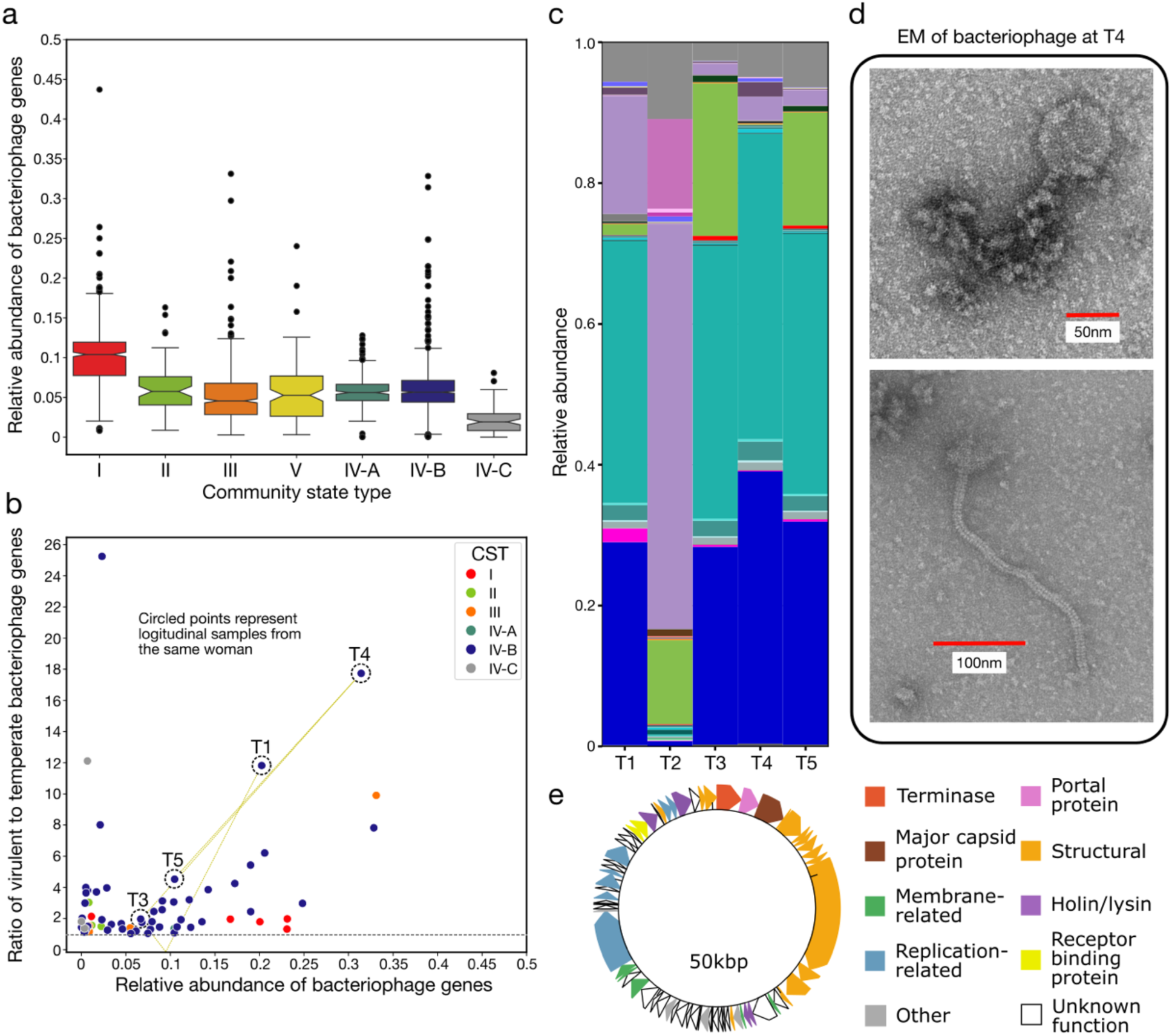
Description of the vaginal phageome as estimated by VIRGO2 and the identification and characterization of a probable lytic bacteriophage bloom. **a,** Relative abundance of bacteriophage genes delineated by the community state type of the corresponding microbiome. **b,** Ratio of the relative abundance of predicted virulent to temperate bacteriophage genes. Points are colored by the community state type of the microbiome and only samples where the relative abundance of virulent bacteriophage surpassed that of temperate bacteriophage are shown. Four of the five metagenomes from a longitudinal collection of samples from the same woman were identified on this plot and are circled by dotted lines and labeled by sequential timepoints separated by ∼2 weeks. **c,** Taxonomic composition of corresponding microbiome at those five timepoints. The taxa color scheme is consistent and can be found in Figs. 2,3,4 and 6. **d,** tunneling electron microscope (TEM) images of bacteriophage-like particles identified in the 0.22µm filtrate of a sample collected from the woman immediately prior to Timepoint 4. TEM images were altered only to increase the size and color of the scale bar. Original images with associated metadata banner can be found in Additional File 1. **e,** Gene annotations of a predicted lytic bacteriophage identified in the assemblies from Timepoints 1,3,4 and 5.

### The landscape of carbohydrate acting enzyme expression differs depending on community composition

A particular strength of VIRGO2 is its ability to analyze and integrate vaginal metagenome and metatranscriptomic data. To demonstrate this feature, we mapped the sequence reads of 344 previously published vaginal metatranscriptomes (see Supplementary Table 6 for accession numbers) to VIRGO2 and then extracted the expression of genes predicted to act on carbohydrates using the available CAZy annotations. The carbohydrate-active expression profiles were then embedded into 2D space using PacMAP^78^, revealing two distinct clusters (Fig. 6a). The first cluster was primarily comprised of microbiomes assigned to CST IV-A, IV-B and II, while the second was comprised of microbiomes assigned to CSTs I, III, and V. We next performed a differential gene expression analysis to identify the specific carbohydrate activities driving the separation of these two clusters. Many of the identified differentially expressed (DE) CAZy functions were predicted to be involved in the catabolism of either glycogen or mucin-associated glycans (Fig. 6b). For example, expression of GHs 20 (β-N-acetylglucosaminidases), 29 (fucosidase), 33 (sialidase), and 101 (endo-α-N-acetylgalactosaminidase) were all higher in samples from cluster 1 whereas GHs 4 (β-glucosidase), 13 (pullulanase), 26 (mannanase), 31 (α-galactosidase), 65 (maltose phosphorylase) were more highly expressed in samples from cluster two (Fig. 6b). The integrated taxonomic and functional annotations of VIRGO2 enabled us to identify the species responsible for the expression of these enzymes. Many of the defining characteristics of cluster 1 were largely expressed by various species of *Prevotella* and *Sneathia* (Fig. 6c), whereas the lactobacilli were mostly responsible the expression of cluster 2 associated enzymes. This result shows that vaginal microbial communities likely metabolize host-produced glycogen but access these resources using different enzymes. However, catabolism of mucin glycans appears to be an activity better accomplished by the microbes that frequent CST IV, due to their expression of enzymes that remove terminal fucose and sialic acid residues.

**Fig. 6.**
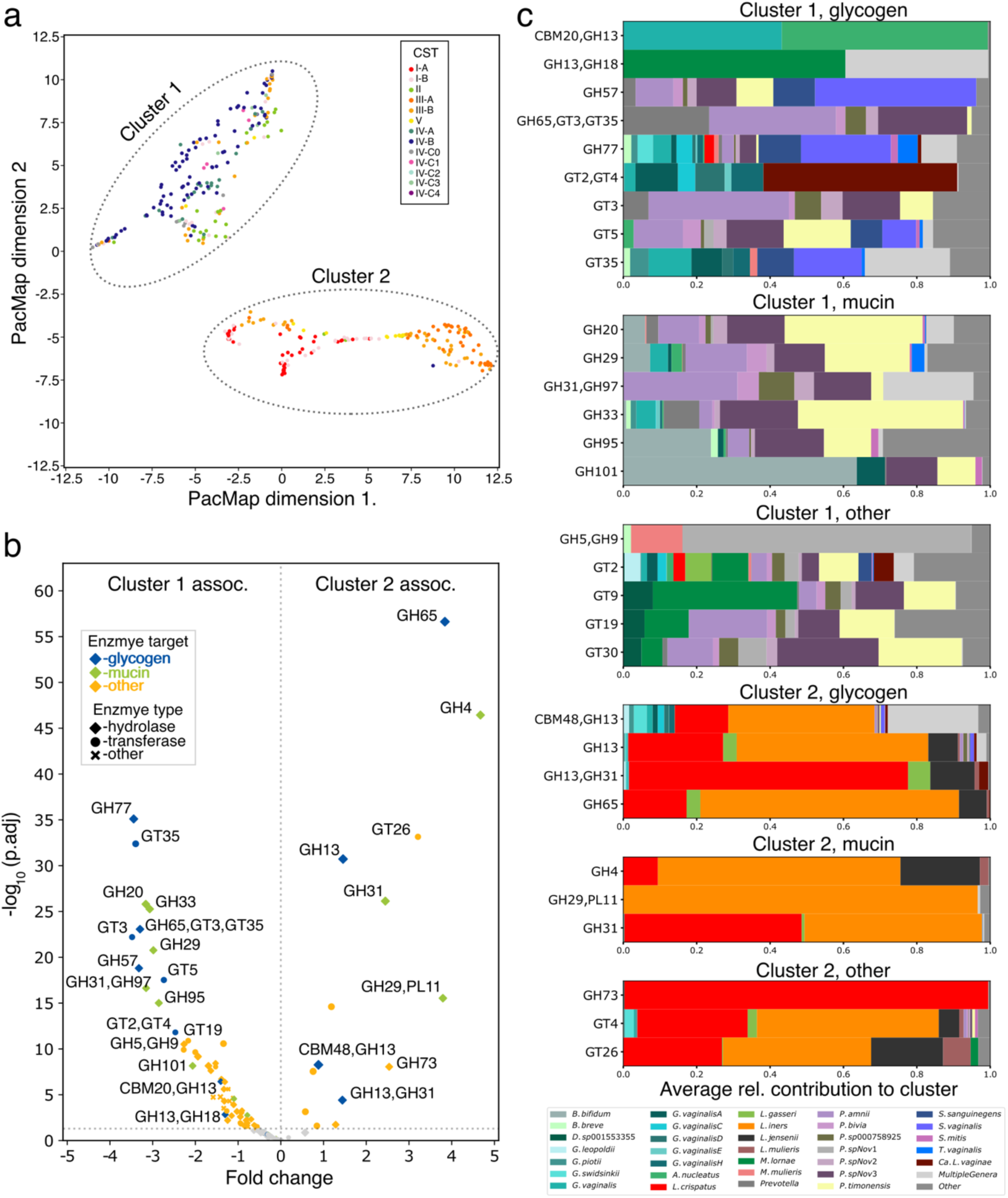
Description of the identification of differences in the expression of carbohydrate-active enzymes by the vaginal microbiome. **a,** Two-dimensional PacMap embedding of the *in vivo* expression patterns of carbohydrate-active enzymes extracted from the VIRGO2 mapping results of 363 vaginal metatranscriptomes. Points are colored by the community state type of the sample. Two clusters were observed. **b,** Volcano plot displaying the results of a differential gene expression analysis comparing clusters 1 and 2. Enzymes are colored by their predicted substrate with a particular emphasis placed on mucin and glycogen. Markers designate the class of the enzyme (hydrolase, transferase, or other). **c,** Taxa driving the expression of significantly differentially expressed carbohydrate-active enzymes in clusters 1 and 2. The displayed bars represent the average contribution of each taxon to the expression of the enzyme in samples from the corresponding cluster.

### *Berryella* sp001552935 is a common yet under-characterized member of the vaginal microbiome

In our initial MAG survey, we found *Berryella* sp001552935 to be an unexpectedly common member of the vaginal microbiome. Subsequent analysis demonstrated that this species had a near universal presence in CSTs IV-A (98.3%) and IV-B (93.0%) communities but rarely comprised more than 10% of these vaginal microbiomes (Fig. 7a). The relative abundance of *Berryella* sp001552935 was also highly diagnostic of CST IV-A and IV-B (receiver-operating characteristic curve, AUC=0.94, Supplementary Fig. 4). To better understand the biology of *Berryella* sp001552935 in the vaginal microbiome, we examined the functions encoded in its genome. The species lacks predicted phosphotransferase systems and appears to be largely asaccharolytic. Acetate and lactate are predicted to enter the cell through a FocA-like porin and then serve as precursors for biosynthetic pathways (Fig. 7b). Energy production, on the other hand, appears to be primarily driven by the import and breakdown of formate and arginine (Fig. 7b). Catabolism of arginine is also predicted to result in the production of the biogenic amine putrescine. The *in vivo* expression of the enzymes in these pathways were confirmed and quantified by extracting the transcriptome of *Berryella* sp001552935 from VIRGO2 mapping results for five metatranscriptomes where the species was well represented (Fig. 7c, minimum 250,000 *Berryella* sp001552935 reads). All enzymatic steps were expressed *in vivo* with average values ranging from 10^2^ to 10^4^ transcripts per million (Fig. 7d). These results give insight into the ecological role of *Berryella* sp001552935 in the vagina and highlight how VIRGO2 can be used to extract and analyze a species’ transcriptome from metatranscriptomic data.

**Fig. 7.**
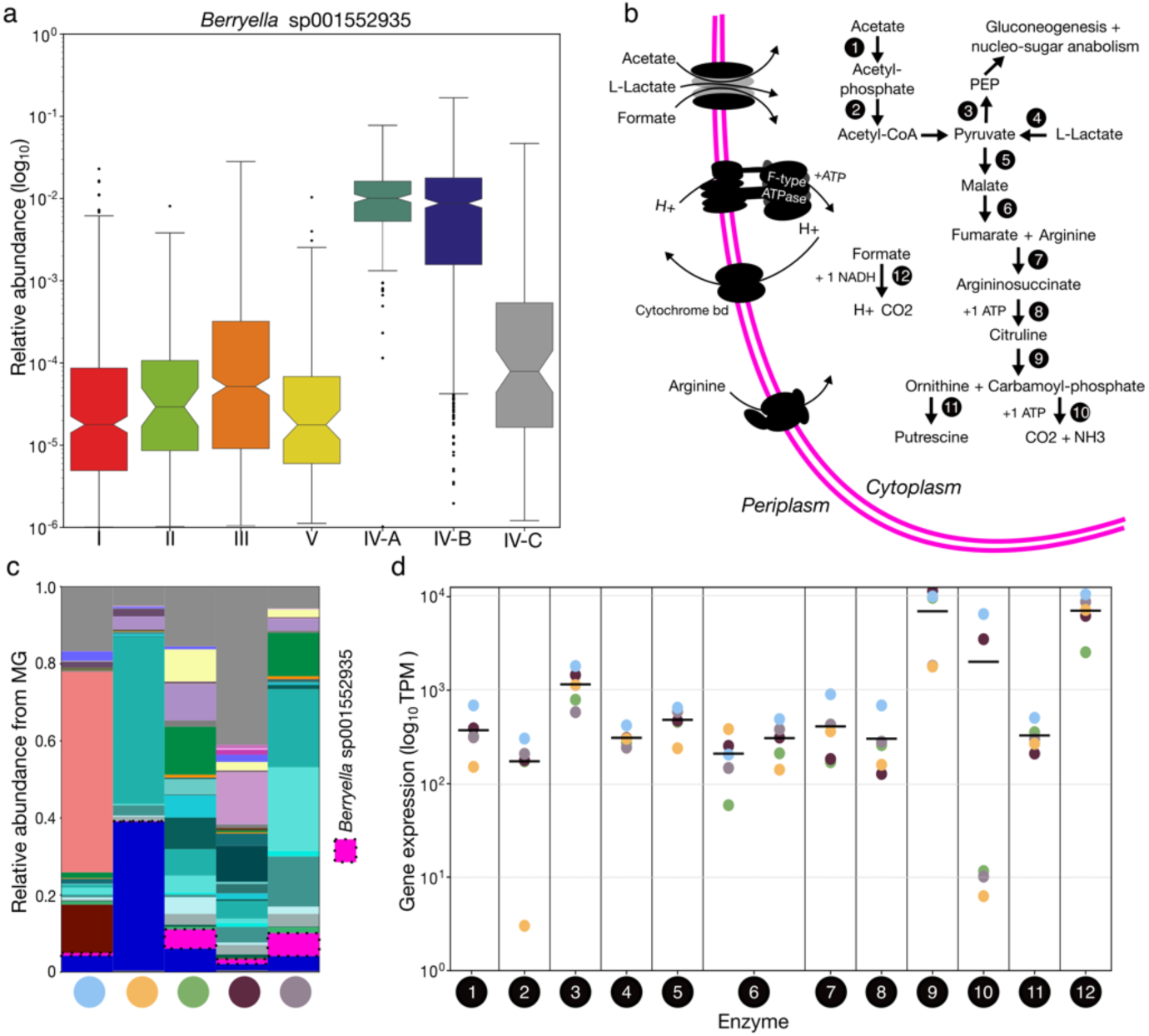
Characterization of the abundance and metabolism of *Berryella* sp001552935 in the vagina. **a,** Log_10_ transformed relative abundance of *Berryella* sp001552935 delineated by the community state type of the sample. **b,** Diagram of predicted metabolic pathways encoded in the genome of *Berryella* sp001552935. Enzyme number designations correspond to: (1) acetate kinase, (2) phosphate acetyltransferase, (3) pyruvate phosphate dikinase, (4) L-lactate dehydrogenase, (5) NAD-dependent malic enzyme (6) hydrolyase, tartrate alpha subunit/fumarate domain protein, Fe-S type, (7) argininosuccinate lyase, (8) argininosuccinate synthase, (9) ornithine carbamoyltransferase, (10) carbamate kinase, (11) diaminopimelate decarboxylase, (12) formate dehydrogenase H. **c,** Relative abundance of *Berryella* sp001552935 in the five metagenomes for which a corresponding metatranscriptome had >250,000 reads mapping in VIRGO2 to the species. The five samples originate from five different North American women and are assigned separate colors, which appear underneath the bars. *Berryella* sp001552935 is highlighted by a dotted outline and all taxa are colored according to the legend found in Figs 2,3,4 and 6. **d,** In *vivo* expression of the enzymes diagramed in subpanel b extracted from the VIRGO2 mapping results for the five metatranscriptomes containing *Berryella* sp001552935. Black bars represent the average expression, and the five points are colored according to the samples characterized in panel **c.**

## Discussion

Genomic databases serve as a critical resource for the robust and reproducible analysis of microbial ‘omics data. Here, we present the construction, validation, and application of VIRGO2, an enhanced iteration of our non-redundant gene catalog for the vaginal microbiome. VIRGO2 was constructed *ab initio* from a set of 2,560 shotgun metagenomes and 3,176 isolate genome sequences and is a complete overhaul of the original database. Sequence data from five continents were used to build VIRGO2, providing a substantial improvement in the database’s geographic representation. VIRGO2 includes twice as many non-redundant genes as VIRGO^20^ and catalogs the genomic diversity of most common and uncommon vaginal bacteria, fungi, and viruses. Over 90% of the genes in VIRGO2 have taxonomic annotations and 80% have at least one functional annotation. The inclusion of data from both MAGs and the residual un-binned contigs also ensures that sequence data from unknown or unidentified organisms are included in the catalog and provides the opportunity for their future annotation. VIRGO2 is also not just a genomic database and its associated annotations. Software to analyze shotgun metagenomes and metatranscriptomes is bundled with the database and was validated to accurately characterize the vaginal microbiome. We anticipate that VIRGO2, and its subsequent releases, will continue to provide a critical resource and tool for the analysis of vaginal microbiome omics data.

A major limitation of the initial VIRGO1 database was the lack of species-level taxonomic resolution within the genus *Gardnerella*. Concurrent with the construction and publication of VIRGO, *G. vaginalis* was split into thirteen species, of which six were given names^64,79^. This reclassification has proven important because diversity within *Gardnerella* has been associated with symptomatic BV^47^ and with preterm birth^27,48,80^. In this iteration of VIRGO, we cataloged the genomic diversity of all described *Gardnerella* species and identified two additional species. Our subsequent analysis reaffirmed that most women with *Gardnerella* in their VMB have several species represented^47,48,80,81^, and demonstrated that the subcommunity of *Gardnerella* spp. has structure^20,47^. This is somewhat unexpected given that these closely related species likely have substantial overlap in their niche use^82^. The vaginal *Lactobacillus* species, which bear a similar degree of relatedness to one another, do not partition their niche space to the same degree as species of *Gardnerella*^33^. It remains to be determined if *Gardnerella* species exhibit mutualistic interactions that drive their co-occurrence^83^, if they engage in resource or spatial niche partitioning^84^, or if they simply cannot exclude one another from the vagina^85^. The limited number of host-produced nutrients (e.g. glycogen, mucin) available to the vaginal microbiome, may preclude resource partitioning as a major driver of *Gardnerella* spp. coexistence. We did find examples of negative correlations between the abundance of *Gardnerella* species, many of which involved *G. swidsinskii*. The genome of this species was found to harbor a unique biosynthetic pathway for the production of a lantibiotic that may be responsible for these negative associations. Future studies, including *in vitro* experimentation, should endeavor to characterize interactions between *Gardnerella* species and how these important bacteria relate to vaginal dysbiosis. Topic models, as employed previously^86^, would naturally lend themselves to the investigation of *Gardnerella* subcommunity structure.

We identified several novel bacterial species in the collection of MAGs used to build VIRGO2. Many of these species were close relatives of known, well described, vaginal bacteria (e.g. novel species in the genera *Gardnerella*) and their biology likely resembles that of their better described sister species, with perhaps some unique aspects. However, *Berryella* sp001552935 stood out as a near-universal member of the CST IV microbiota that is not often discussed in the literature. The species is in the family *Eggerthellaceae* and may have been referred to as “*Eggerthella* sp.” or “*Eggerthella* type 1” in a few 16S rRNA gene sequencing-based studies^25,87–91^. Twice this “Eggerthella type 1” species has been associated with all four clinical signs of the Amsel criteria for diagnosis of bacterial vaginosis^88,89^, concordant with our finding of *Berryella* sp001552935’s diagnostic value for CST IV. Due to the species’ specificity for non-*Lactobacillus* dominant microbiota, it is difficult to disentangle its association with vaginal health from that of other BV-associated microbes. Nevertheless, one study did find “*Eggerthella* type 1” to be correlated with higher IL-1⍺, IL-1β, and TNF-⍺, and two found it to be associated with increased risk of HIV acquisition^25,91^. Based on its genomic contents and *in vivo* transcriptional activity, we predict *Berryella* sp001552935 to be largely asaccharolytic, like its relative *E. lenta*^92^, and to similarly rely on acetate and arginine as sources of carbon and energy. Acetate is the end-product of the fermentative activities of several CST IV community members^93^ and is one of the most common molecules in the vagina when lactobacilli are not present^39^. Arginine is also a common component of vaginal fluid and can be found at high concentrations in seminal fluid^94^. The catabolism of arginine by *Berryella* sp001552935 is predicted to result in the production of the biogenic amine putrescine, which can function to raise vaginal pH and likely contributes to vaginal malodor. We therefore posit that *Berryella* sp001552935 acts as a decomposer in the vaginal ecosystem, upcycling waste products into biomass and energy and aiding the growth of other BV-associated microbes.

We placed an emphasis on cataloging the non-bacterial components of the vaginal microbiome in VIRGO2 to expand the utility of the database. Several fungal species were identified in the shotgun metagenomics data used to build VIRGO2 and more were added through the inclusion of isolate whole genome sequences. Our analysis of the shotgun metagenomes revealed that *C. albicans*, *N. glabratus* and *P. kudriavzevii* were the primary components of the vaginal mycobiome, concordant with past survey data^95–97^. Some studies have identified other species as minor components of the fungal communities^98,99^; however, we found no evidence of these species in the shotgun metagenomes. It could be that they are present but only below the limit of detection of metagenomics (<0.01% relative abundance), or they could simply represent fungal contaminants from the air or molecular reagents^100^. We also found that only 13.2% contained a detectable abundance of fungi, a rate similar to past microscopy-derived estimates of asymptomatic *Candida* carriage^101,102^. These fungi containing metagenomes were more likely to have a bacterial community that was dominated by *L. iners,* than any other VMB compositions. It is likely that detection of fungi in metagenomic data is harder in CST IV bacterial communities due to their higher population sizes^103^. Past observational studies have also found associations between *L. crispatus*^104,105^ or *L. iners*^43^ and detection of vaginal yeast. Symptomatic vulvovaginal candidiasis (VVC), on the other hand, has been repeatedly linked to BV and BV indicated antibiotic therapy^106^. It is commonly thought that lactobacilli temper the virulence of vaginal fungi, although the exact mechanism has yet to be elucidated. The inclusion of fungal sequence data in VIRGO2 enables the characterization of the vaginal mycobiome using metagenomic and metatranscriptomic data and will advance our understanding of its relationship to the bacterial microbiome.

Our efforts to include, identify, and catalog viral sequences in VIRGO2 enable its use for the characterizing the vaginal virome. We found that bacteriophages were responsible for the majority of the virome, with eukaryotic viruses (e.g. human papillomavirus) representing only a minor fraction, consistent with past studies^107,108^. Bacteriophage have long been theorized to be major drivers of compositional changes in vaginal bacterial communities^109^. Many women have *Lactobacillus*-dominant bacterial communities; a virulent bacteriophage capable of infecting the dominant *Lactobacillus* could collapse its population, resulting in a dramatic shift in community composition. Yet, we found that most bacteriophage in the vagina are predicted to have temperate lifestyles. A high abundance of virulent bacteriophages was observed in only a handful of metagenomes, and the bacterial communities in these cases tended not to be dominated by lactobacilli. These results suggest that virulent bacteriophage are not major drivers of bacterial community composition and are likely not responsible for most transitions away from *Lactobacillus* dominance. Temperate bacteriophage integrate into their host bacterium’s genome and excise themselves in response to a variety of environmental or metabolic triggers^110^. Vaginal *Lactobacillus* are known to harbor numerous prophage in their genomes^111–113^, but the conditions that induce their excision are not well characterized^114^. Induction of temperate bacteriophage could impact community composition by reducing the population size of their host^115^; however, prophage can also function to protect their host from infection by other bacteriophage^116^, regulate host gene expression^117^, kill closely related competitors^118^, regulate population density^119^, or act as agents of horizontal gene transfer^120^. Going forward, VIRGO2 will facilitate the characterization of the temperate bacteriophage common to the VMB and how they influence community dynamics and vaginal health.

To demonstrate how VIRGO2 can be used to analyzed vaginal metatranscriptomes, we examined the expression of carbohydrate acting enzymes in a previously published dataset^121^. The results from this analysis identified several CAZy families that were differentially expressed between communities that were dominated by *Lactobacillus* versus those that were not. Many of these differentially expressed enzyme families were predicted to be involved in the breakdown of either glycogen or mucin-associated glycans. This result suggests that, while vaginal bacteria have substantial overlap in their resource use, the enzymatic functions they use to access these resources, and the extent to which they utilize them, are different. For example, communities containing *Prevotella* and *Gardnerella* spp. might be more capable of catabolizing mucin glycan chains because they express the enzymes necessary to remove terminal sialic acid and fucose residues, providing access to the mucin-associated glycans^50,122^. The vaginal lactobacilli do not express these enzymes and therefore may only have limited access to those mucin glycan chains. Glycogen, on the other hand, appears to be a major source of carbon and energy for most vaginal bacteria but is catabolized using slightly different enzymatic functions. Lactobacilli primarily use GH13 and GH33 domain containing enzymes, whereas many of the bacteria that comprise CST IV communities (e.g. *Gardnerella* spp., *Prevotella* spp., or *Sneathia* spp.) express GH77 and GT35 domain containing enzymes. These differences could be harnessed to selectively promote the growth of lactobacilli or inhibit the growth of non-optimal bacteria. For example, a recent study by Pelayo et al. identified several small molecule inhibitors of *Prevotella* and *Gardnerella* sialidases that could limit the growth of these bacteria in the vagina^122^. Another study found that supplementation with maltose, a metabolite of glycogen produced by vaginal lactobacilli^123^, was able to shift the macaque vaginal microbiota to *Lactobacillus-*dominance^124^. Additional studies aimed at understanding microbial resource use in the vagina, including beyond carbohydrate metabolism characterized here, could reveal additional means to modulate community composition.

Integrative multi-omic analyses can provide mechanistic insight into the interactions between humans and their resident microbes. Yet a major barrier to this discovery is the lack of computational resources for the analysis of these data. VIRGO2 helps fill this critical need by providing an analytical framework for the analysis and integration of vaginal metagenomic and metatranscriptomic data. Here, we present a major update to this resource that 1) expands the geographic coverage of the constituent sequence data, 2) broadens the scope of the database to include non-bacterial microbes including fungi and viruses, and 3) remasters the taxonomic and functional annotations to match current standards. Still, VIRGO2 is not without limitations. Although sequence data from five continents were included in its construction, most continents are only represented by a single dataset from a single country, and South American populations are not represented. The database is also missing representation of pre- and post-reproductive age individuals as no shotgun metagenomes were available for these populations. Despite these limitations, we expect VIRGO2 to enhance the analysis of vaginal microbiome ‘omics data to better understand how these not-so-simple microbial communities impact health.

## Acknowledgements

Research reported in this publication was supported in part the National Institute of Allergy and Infectious Diseases (NIAID), the National Institute for Nursing Research (NINR) of the National Institutes of Health (NIH) under awards UH2AI083264, R01AI116799, U19AI084044, R01NR014784 and R01NR015495, and from the Gates Foundation under awards OPP1189217, INV048956, and INV-048982. I. C. was supported by NIAID training grant T32AI162579. D.A.R. was supported by the Thomas C. and Joan M. Merigan Endowment at Stanford University. We thank the UMSOM Electron Microscopy Core Imagining Facility for assistance with electron microscopy. We also thank members of the Vaginal Microbiome Research Consortium (VMRC) for helpful discussions.

## Author Contributions

M. T. France, Conceptualization, Methodology, Software, Validation, Data curation, Formal analysis, Writing – original draft, Writing – review and editing | I. Chaudry, Software, Formal analysis, Validation, Writing – review and editing | L. Rutt, Investigation, Data curation, Supervision, Writing – review and editing | M. Quain, Investigation, Writing – review and editing | B. Shirtliff, Investigation, Writing – review and editing | E. McComb, Investigation, Writing – review and editing |F. A. Hussain, Formal analysis, Writing – review and editing | A. Maros, Formal analysis, Writing – review and editing | M. Alizadeh, Formal analysis, Writing – review and editing | MA. Elovitz, Funding Acquisition, Supervision, Writing – review and editing | D. Relman, Funding Acquisition, Supervision, Writing – review and editing | A. Rahman, Funding Acquisition, Supervision, Writing – review and editing | R. M. Brotman, Funding Acquisition, Supervision, Writing – review and editing | J. Price, Funding Acquisition, Supervision, Writing – review and editing | M. Kassaro, Funding Acquisition, Supervision, Writing – review and editing | J. Holm, Formal Analysis, Supervision, Writing – review and editing, | B. Ma, Conceptualization, Writing – review and editing | Jacques Ravel, Conceptualization, Funding acquisition, Supervision, Writing – review and editing. All authors contributed and approved the final version of the paper.

## Resource availability

### Lead contact

Additional information and requests for additional data and code should be directed to and will be fulfilled by the Lead Contact, Jacques Ravel

### Data and code availability

The VIRGO2 software, the non-redundant nucleotide gene catalog and database, the VOG amino acid protein family database, the associated annotation files, and detailed tutorials are available at: http://virgo.igs.umaryland.edu. The metagenomes used to construct VIRGO2 are all publicly available can be obtained from either the National Center for Biotechnology Information (NCBI) Sequence Read Archive (SRA) or from the European Nucleotide Archive using the accession numbers provided in Supplementary Table 1. All scripts used in the described analyses and to construct the displayed figures are available at: https://github.com/ravel-lab/VIRGO2.

## Methods

### Generation of metagenome assembled genomes

A total of 2,560 shotgun metagenomes were used in the generation of the VIRGO2 database (Supplementary Table 1). The vast majority of these metagenomes were generated from human vaginal swab or lavage specimens (n=2503, ∼98%). However, a small proportion were generated from male urethral swab specimens, which were generated as part of a sexual network study (n=57, ∼2%). Sequence reads originating from the human genome were identified and removed in all shotgun metagenomes using BMTagger (ftp://ftp.ncbi.nlm.nih.gov/pub/agarwala/bmtagger, default settings). This included metagenomic datasets downloaded from public databases to ensure all host reads were removed prior to processing. The host-removed datasets were further processed using fastp^125^ to remove barcode sequences and trim low-quality bases (v0.21.0, -l 50 - W 4 -M 20). The median number of reads per metagenome post-processing was 1.27×10^7^ (range from 2.0×10^4^ to 3.3×10^8^) and only 3.4% of metagenomes had fewer than 10^6^ reads. Metagenomes originating from studies performed in Fiji, Ireland, and China were not sequenced to the same depth as the rest of the metagenomes and therefore had fewer reads per sample on average (median: 1.6×10^6^; range: 2.0×10^4^-4.8×10^7^). The metagenomes sequenced from male urethral swabs also had a greater proportion of host-reads resulting in fewer available microbial reads (median: 1.4×10^6^, range: 2.8×10^5^-6.3×10^7^). *De novo* assembly was attempted on all shotgun metagenomes using metaspades^126^ (v3.15.3, default settings) and 2,548 (99.5%) produced assemblies. The raw assemblies were then separated into a metagenome assembled genomes (MAGs) using a custom multipronged approach whereby binning was performed on each metagenome using MaxBin2^127^ (v2.2.7, -min_contig_length 500), metaBat2^128^ (v2.12.1, -- maxEdges 500), and Metadecoder^129^ (v1.0.8, default settings). The resulting set of bins were then compiled using metaWRAP^130^, which relies on checkM^131^ to estimate MAG completeness and degree of contamination. All MAGs were further processed using MAGpurify^132^ (v2.1.2, default workflow) and cleaned versions were accepted where the degree of contamination, as estimated using checkM, decreased, without a concomitant decline in estimated completeness >1%. MAGs demonstrating a final completion level >70% and <10% contamination were used in the analysis and in the construction of the VIRGO2 database (n=15,142, Supplementary Table 2). A total of 21,202,535 genes were then identified on the MAGs using Prodigal^133^ (v2.6.3, -c -p single).

### Processing residual metagenome assembly contigs

Contigs greater than 2,000 bp that were not incorporated into any bin, were processed as “un-binned contigs”. These contigs likely represent either fragments of bacterial genomes which were not successfully binned or fragments of non-bacterial genomes (e.g. fungi, bacteriophage). A total of 8,294,162 genes were identified on the un-binned contigs using Prodigal^133^ (v2.6.3; -c -p meta).

### Cultured isolate genome (CIG) sequences

Whole genome sequences of known vaginal bacteria, fungi, and viruses were used to supplement the metagenomic data in the construction of the VIRGO2 catalog. A total of 4,013 genomes were downloaded representing 198 species, of which 190 were bacteria, 4 were fungi (*Candida albicans*, *Candida dubliniensis*, *Nakaseomyces glabratus*, and *Pichia kudriavzevii*), 3 were viruses (human immunodeficiency virus, human papilloma virus, and human alpha herpes virus), and 1 was a protist (*Trichomonas vaginalis*). A complete list of the species targeted, and accession numbers for these genome sequences can be found in Supplementary Table 3. Sequences were acquired from NCBI in February of 2023. A total of 8,015,873 genes were identified on the CIGs using Prodigal^133^ (v2.6.3, -c -p single).

### Generation of non-redundant gene clusters

All genes identified in the three data sources (MAGs: 21,202,535, un-binned contigs: 8,294,162, CIGs: 8,015,873) were clustered using CDHIT^134^ (v4.8.1, cd-hit-est -n 8 -c 0.95 -aS 0.9) with thresholds of 95% sequence identity, and 90% overlap of the shorter sequence. This process produced 1,880,579 gene clusters, each represented by their longest sequence constituent and hereafter referred to as “non-redundant genes”. The median number of sequences per cluster was 2 (range 1-6,464), and 729,189 clusters (38.8%) were comprised of single sequences.

### Identifying and removing anomalous non-redundant genes

Non-redundant genes were searched against the human genome^135^ (NC_060925.1, T2T- CHM13v2.0) and against the SARS-CoV-2 genome^136^ (NC_045512.2) using blastn^137^ (v2.10.0, - evalue 0.00001) to identify (percentage identity ≥75%, coverage ≥ 50%) and remove these undesired sequences from the VIRGO2 database. A total of 34,321 non-redundant genes were removed due to matches to the human genome and 20 due to matches to the SARS-CoV-2 genome. An additional 9,159 non-redundant genes which contained at least one “N” in their sequence were removed. An all-versus-all blastn (v2.10.0, -evalue 0.00001) search was then performed against the remaining non-redundant genes to identify those which contain a stretch of sequence, of at least 100bp, identical to that found in another non-redundant gene. For these cases of exact contiguous matches, only the longer gene was kept, resulting in the removal of an additional 37,665 genes. A final filtering step was applied to identify non-redundant genes whose prevalence, abundance, and tetramer content were dissimilar from that of the overall species’ distribution. These genes were primarily repeats of adapter or barcode sequence and were removed from the database (n=26,897). The final number of VIRGO2 non-redundant genes following these filtering steps was 1,772,517.

### Assigning taxonomy to VIRGO2 non-redundant genes

The taxonomy of the VIRGO2 source sequences (MAGs, CIGs, un-binned contigs) form the basis of the annotations applied to the non-redundant gene clusters. Taxonomy was first assigned to the MAGs and CIGs using GTDB-tk^138^ (v2.1.1, GTDB release 207_v2, classify_wf). Then, a two-stage strategy was used to assign taxonomy to the un-binned contigs. First, the un-binned contigs were searched against the NCBI nt database (accessed March 2023) using blastn (v2.10.0, -evalue 0.00001). Second, the un-binned contigs were searched, using the same settings, against a blast database constructed from the MAGs and CIGs. Taxonomic annotations were propagated to the unbinned contigs preferentially from the MAGs and CIGs over those from matches to the NCBI nt database. The taxonomic annotations from the source sequences were then applied to the final non-redundant gene clusters based on the taxonomy of their constituents using the following rules: (1) gene clusters with uniform species-level taxonomy were assigned to that singular species (n=1,193,357); (2) gene clusters with variable species-level taxonomy, but a majority (>50%) of one, were assigned to that species (n=279,714); (3) gene clusters without a majority species-level taxonomy but a majority (>50%) genus-level taxonomy, were assigned to that genus (n=71,195); (4) genes without a majority genus-level taxonomy were assigned as “MultiGenera” genes (n=68,548); (5) genes without any available taxonomic information were not assigned any taxonomic annotation (n=159,703).

### Assigning function to VIRGO2 non-redundant genes

A rich suite of functional annotation schemes was applied to the non-redundant genes. EggNOG-mapper^65^ (v2.1.10, eggNOG database v5.0.2, -m diamond, -evalue=0.001) was used to annotate the genes with the following schemes: eggNog^68^, COG^139^, PFAM^70^, GO terms^71,140^, EC numbers^141^, BiGG Reaction^73^, KEGG^74^, and Carbohydrate-active enzymes^75^ (CAZy). Antibiotic resistance genes and virulence factors were identified and annotated using AMRFinderPlus^67^ (v3.10.16, database v2022-11-2, default settings). Secondary metabolite biosynthetic gene clusters were identified in the MAGs using antiSmash (v7.1.0, --genefinding-tool prodigal, --clusterhmmer, -cc-mibig, --cb-general, --cb-subclusters, --cb-knownclusters) and then the annotations were propagated to the appropriate NGR cluster^76^. The VIRGO2 non-redundant genes were translated into protein sequences using biopython (v1.81) and the appropriate codon table. Orthologous protein sequences were identified for each species, individually, with OrthoFinder^142^ (v2.5.4, default settings). The 1,772,517 non-redundant genes were reduced to 1,032,117 orthologs.

### Identifying and annotating bacteriophage sequences in VIRGO2

Bacteriophage sequences among the raw metagenome assemblies were identified using virSorter2^143^ (v2.2.4, --min-length 3000 --min-score 0.5) and their quality was assessed using checkV^144^ (v1.01, database v1.5, default settings). Predicted bacteriophage sequences were only used when they met the following conditions: (1) more than one known viral gene detected, (2) sequence not predicted to be a prophage, (3) fewer than 10 bacterial host genes detected, and (4) checkV completeness estimate > 90%. This conservative filtering process produced 3,414 predicted bacteriophage genome sequences which encoded 43,365 of the VIRGO2 non-redundant genes. Bacteriophage lifestyles (lysogenic versus virulent) were predicted using PhaTYP^145^ (v1, default settings) and the sequences were classified at the family level using PhaGCN^146^ (v1, default settings). Bacteriophage host identity was predicted using Prokaryotic virus Host Predictor^147^ (PHP, v1, default settings) with a dereplicated set of the VIRGO2 MAGs (dereplication performed using dRep^148^, v3.1.1, --sa 0.975, --S_algorithm fastANI, --clusterAlg ward) used as the host database. PHP host predictions were accepted when the reported score was ≥ 1400 and were recorded at the genus level. Bacteriophage taxonomy, host, and lifestyle annotations were propagated to the appropriate non-redundant genes (Supplementary Table 5).

### Using VIRGO2 to profile *In silico* generated mock communities, vaginal shotgun metagenomes and, metatranscriptomes

Software to map reads to the VIRGO2 non-redundant gene catalog and to characterize metagenome/metatranscriptome composition is provided along with the catalog and is available online (virgo.igs.umaryland.edu). Briefly, reads are mapped to the non-redundant gene catalog using Bowtie2 (v4.1.2, -N 0,-L 20 -D 20,-R 3,-k 10,--sam-no-qname-trunc,--end-to-end,--seed 343, --no-unal) and the resulting sam file is parsed using samtools (v1.11, sort, view commands) and custom python3 code. To validate VIRGO2’s ability to characterize vaginal communities, we generated mock shotgun metagenomes using inSilicoSeq (v1.5.3, --seed 343, -n 10m, --model novaseq)^149^. Isolate genome sequences of several vaginal bacteria were used to generate mock shotgun metagenomes comprising known abundances of species. The exact isolates used are denoted in Supplementary Table 3. Accuracy of the resulting quantifications was assessed using a linear regression of observed versus expected relative abundances of bacteria using the scipy python package (v1.1.10). Following these validation efforts, the 2,560 metagenomes used to construct VIRGO2 and as well as 363 vaginal metatranscriptomes (Supplementary Table 1) were mapped to the catalog using the same approach. The results of these mappings were used in the subsequent analyses generating Figures 4-7 (metagenomes: Figures 4, 5, and 7; metatranscriptomes: Figures 6 and 7). Relative abundances were quantified using the sum of gene coverages for each bacterium, fungi, or virus.

### Identifying correlations in the abundance of *Gardnerella* spp

The number of *Gardnerella* species identified in each metagenome was determined using a conservative 1% relative abundance threshold for detection. Correlations in the relative abundance of *Gardnerella* species were quantified using ANCOMBC (v2.0.3, secom_linear, method=pearson, thresh_hard=0.3)^150,151^. A lantibiotic operon was identified in the genome of *G. swidsinskii* C0102A2 using BAGEL4 (v.1.2, default settings)^152^. The lantibiotic gene was then identified in the genomes and MAGs of other *G. swidsinskii* using blast searches (blastp, v2.13.0, default settings)^137^.

### Characterization of vaginal bacteriophage

Genes annotated as originating from bacteriophage were identified using the appropriate annotation table included in VIRGO2 (10.VIRGO2.phage.txt, available at: virgo.igs.umaryland.edu). The overall relative abundance of bacteriophage was quantified using the sum of gene coverages for the bacteriophage genes divided by the sum of coverages for all genes. To quantify the ratio of virulent to temperate bacteriophage, the sum of coverages of the bacteriophage genes annotated as virulent was divided by the sum of coverage of those annotated as temperate. Four of the 63 metagenomes with higher coverage of virulent than temperate genes were identified as originating from the longitudinal sampling of a single person. The high coverage of virulent bacteriophage genes was traced back to a single high coverage bacteriophage contig. The contig’s coding sequences were identified and annotated with Pharokka (v1.7.3, -g prodigal-gv, --dnaapler)^153^. Bacteriophage was isolated from a vaginal swab specimen collected one day prior to fourth timepoint, that had been stored at −80℃ without any preservation media^53^. The sample was eluted off the swab by vortex in 2 mL of sterile phosphate-buffered saline (PBS) for 2 min. The elution was sequentially filtered through a 5 µm, 0.45 µm, and 0.22 µm diameter filters (Thermo Fisher Scientific, Waltham, MA, USA). The final filtrate was used to prepare a negative stain TEM grid at the University of Maryland School of Medicine (UMSOM) Electron Microscopy Core. A formvar/carbon coated TEM grid underwent plasma cleaning (glow discharge) before 5 µL of the eluted bacteriophage sample was applied to formvar side of the grid. The sample was given 1 min to adsorb to the grid, and excess sample was removed. Next, a fixative (4% glutaraldehyde in 0.12 M sodium cacodylate buffer, pH = 7.2) was added to the grid for 5 min, and then washed four times with deionized water. Lastly, the grid was stained with 1% aqueous uranyl acetate for 1 min. Grids were dried at room temperature overnight before mounted on the TEM. Micrographs were collected on the FEI tecnai T12 instrument (Field Electron and Ion Company, Hillsboro, OR, USA).

### Analysis of carbohydrate-active enzyme expression

The transcriptional activity of the analyzed 344 vaginal metatranscriptomes was quantified as transcripts per million (TPM). The expression of genes encoding carbohydrate-active enzymes^75^ were identified and extracted using the VIRGO2 CAZy annotation table (8.VIRGO2.CAZy.txt, virgo.igs.umaryland.edu). The resulting dataset of CAZy enzyme expression was processed using the Pairwise Controlled Manifold Approximation (PacMap) dimensionality reduction method (v0.7.0, n_components=2, MN_ratio=0.5, FP_ratio=2.0, distance=angular), revealing two clusters. Differential expression analysis was conducted using VariancePartition (v1.28.9, default settings)^154^ between the two groups to identify the CAZy enzyme classes responsible for the clustering.

### Characterization of *Berryella* sp001552935

The relative abundance of *Berryella* sp001552935 in samples from different CSTs was quantified as described above. Metabolic pathways were outlined via the KEGG orthologs assigned to a representative *Berryella* sp001552935 MAG (MAG14846, Supplementary Table 2) by BlastKOALA (v3.1, prokaryotes)^155^ and then matched to VIRGO2 genes via blastp (blastp, v2.13.0, default settings)^137^. Five of the 363 vaginal metatranscriptomes were identified as having more than 100,000 reads mapping to *Berryella* sp001552935 genes. These samples were used to estimate the expression of genes in the putative metabolic pathways used by the bacterium to generate energy and assimilate carbon in the vagina.

## Supplementary Table Legends

**Supplementary Table 1: Accession numbers of metagenomes included in VIRGO2**

**Supplementary Table 2: Inventory of metagenome assembled genomes generated in this study**

**Supplementary Table 3: Accession numbers of isolate whole genome sequences included in VIRGO2**

**Supplementary Table 4: Prevalence of vaginal bacteria across different geographies**

**Supplementary Table 5: Inventory of bacteriophage genome sequences identified in VIRGO2**

**Supplementary Table 6: Accession numbers of vaginal metatranscriptomes used in our analyses**

## Supplementary Figure Legends

**Supplementary Figure S1:** Histogram displaying the number of non-redundant genes by the size of the cluster

**Supplementary Figure S2:** Percent of VIRGO2 non-redundant genes assigned to various functional annotation schemes

**Supplementary Figure S3:** Metagenome accumulation curves of prominent vaginal bacteria assessed at the level of orthologs (VOGs)

**Supplementary Figure S4:** Receiver operating characteristic curves demonstrating the ability of the relative abundances of select microbial species (*Berryella* sp001552935, *G. vaginalis*, *G. swidsinskii*, *P. timonensis*, *F. vaginae*, *M. lornae*, and *A. indicium*) to predict an overall community composition assigned to CST IV-A or IV-B.

**Additional File 1:** Fasta file containing the coding sequences of the Lantibiotic operon identified in *G. swidsinskii*

**Additional File 2:** Original, unaltered TEM micrographs used in Figure 5.

