## Supplementary figures and images for "VIRGO2: Unveiling the Functional and Ecological Complexity of the Vaginal Microbiome with an Enhanced Non-Redundant Gene Catalog"

### Supplementary Figure 1

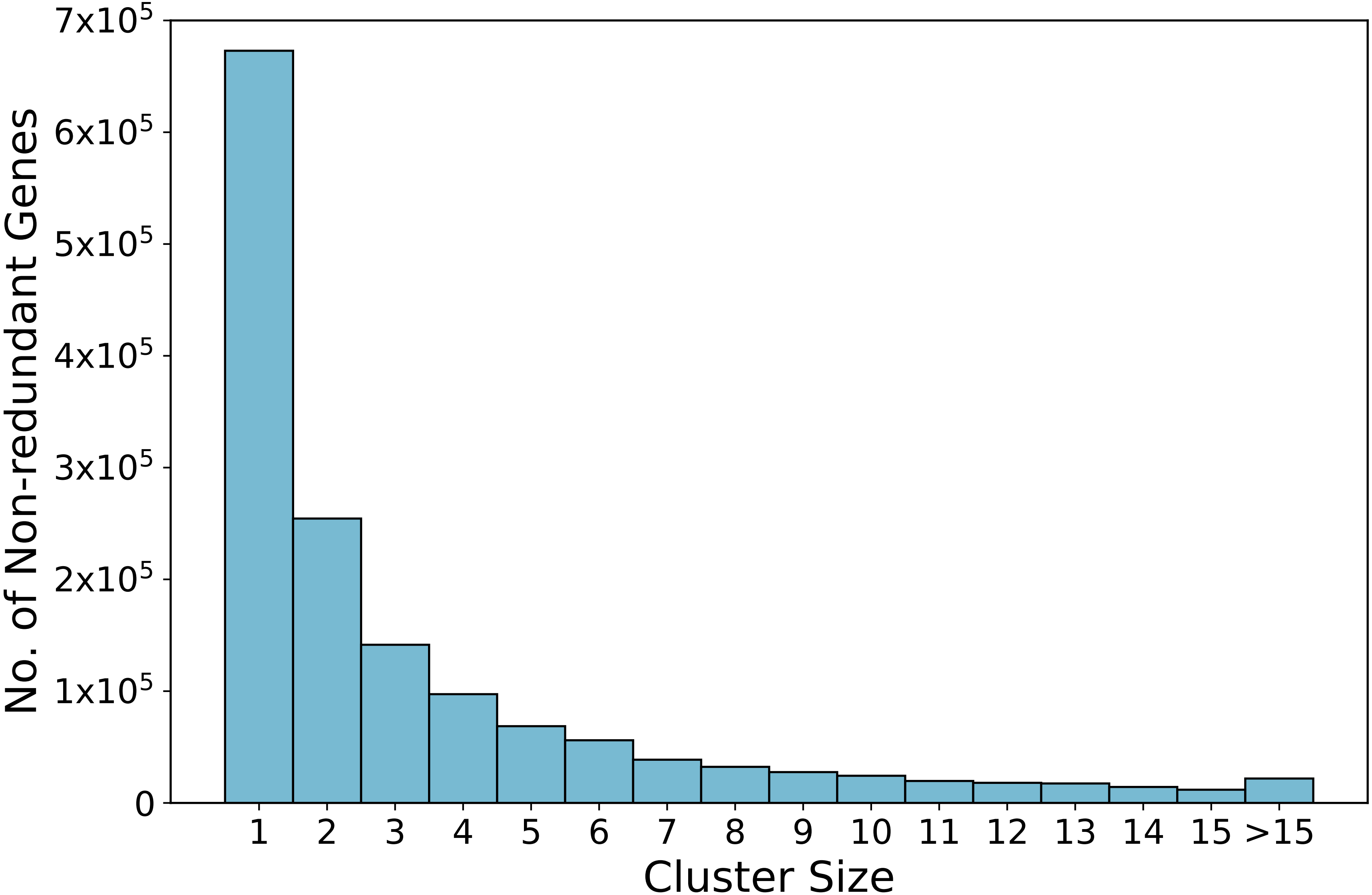

### Supplementary Figure 2

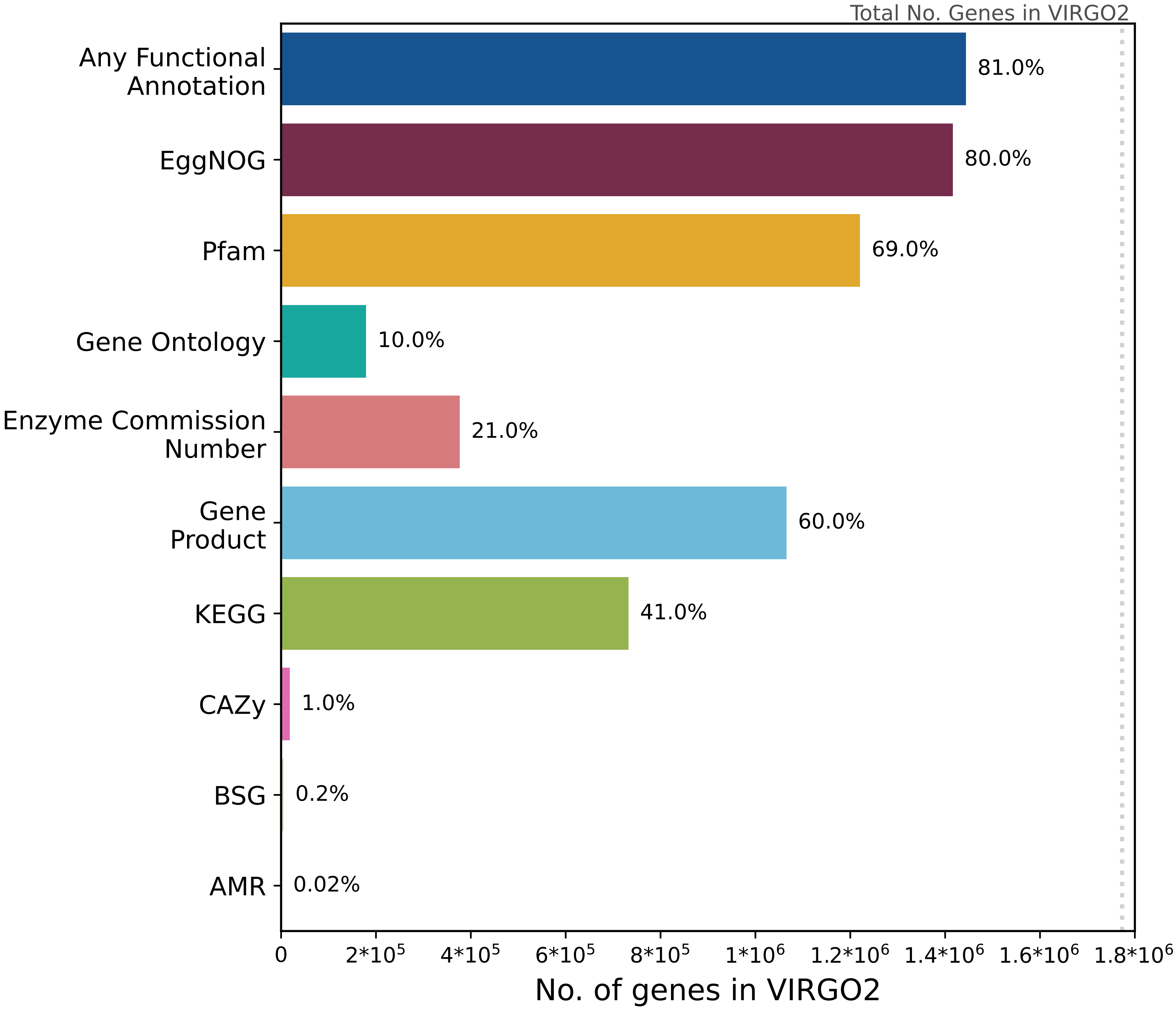

### Supplementary Figure 3

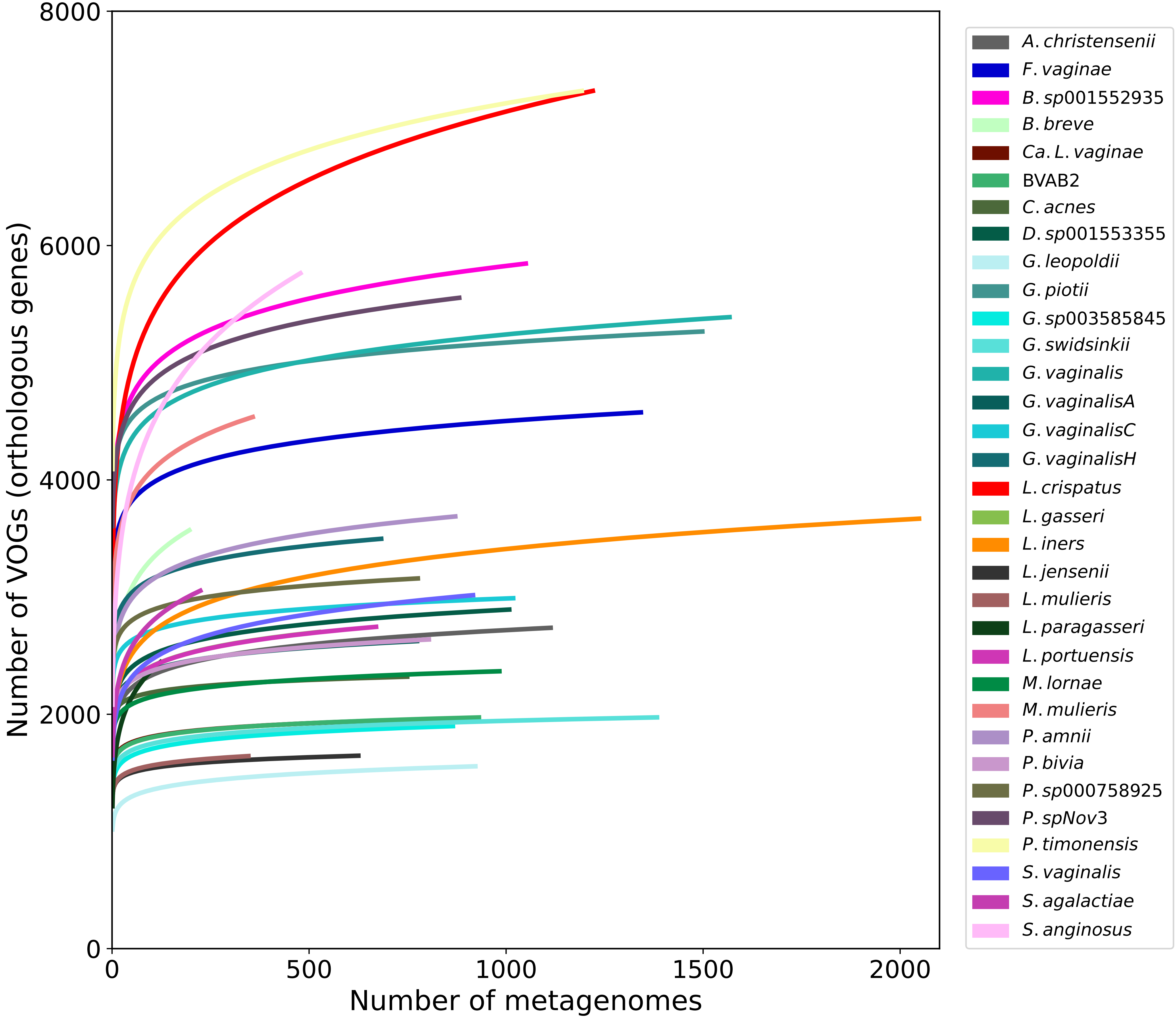

### Supplementary Figure 4

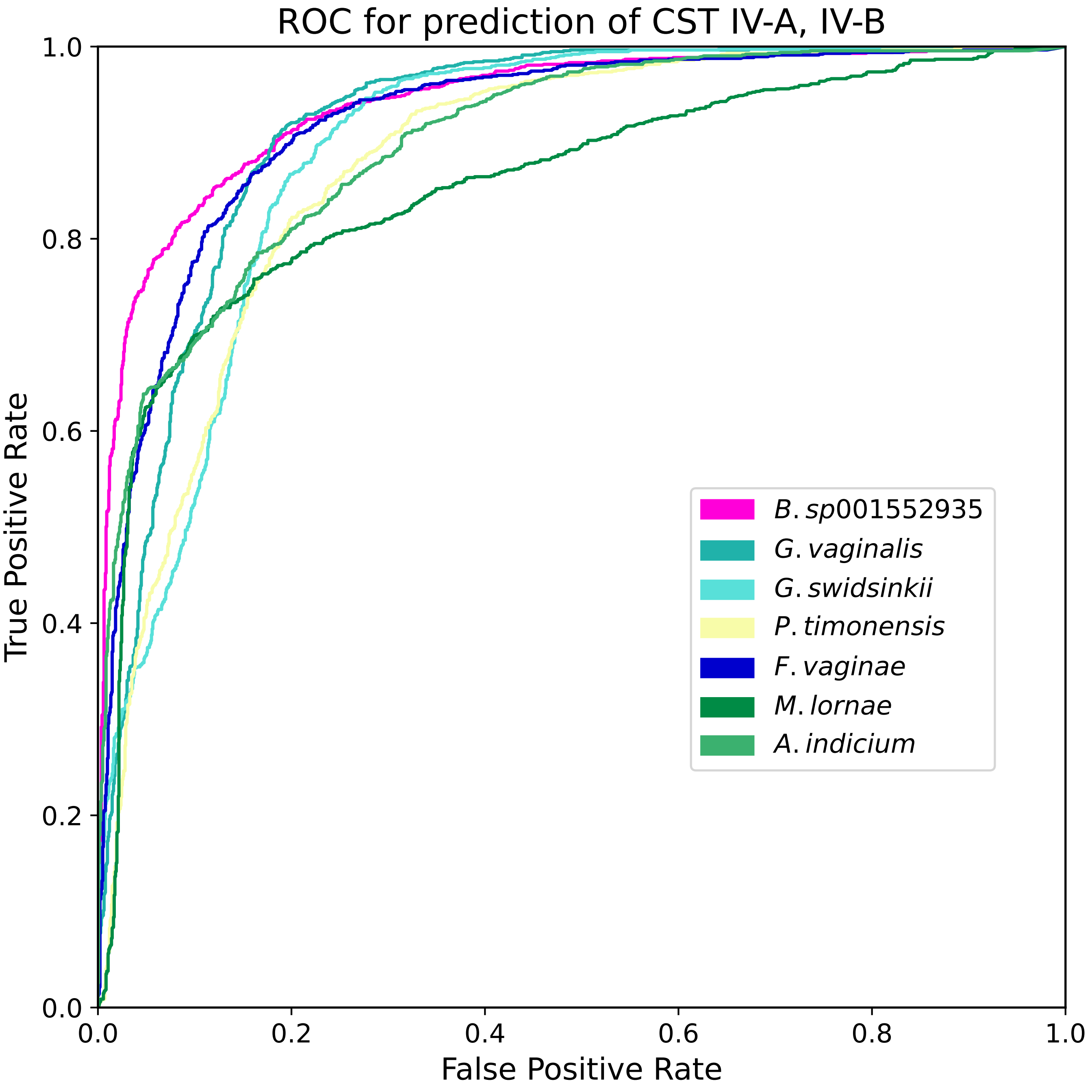
