## Additional File 2 for "VIRGO2: Unveiling the Functional and Ecological Complexity of the Vaginal Microbiome with an Enhanced Non-Redundant Gene Catalog"

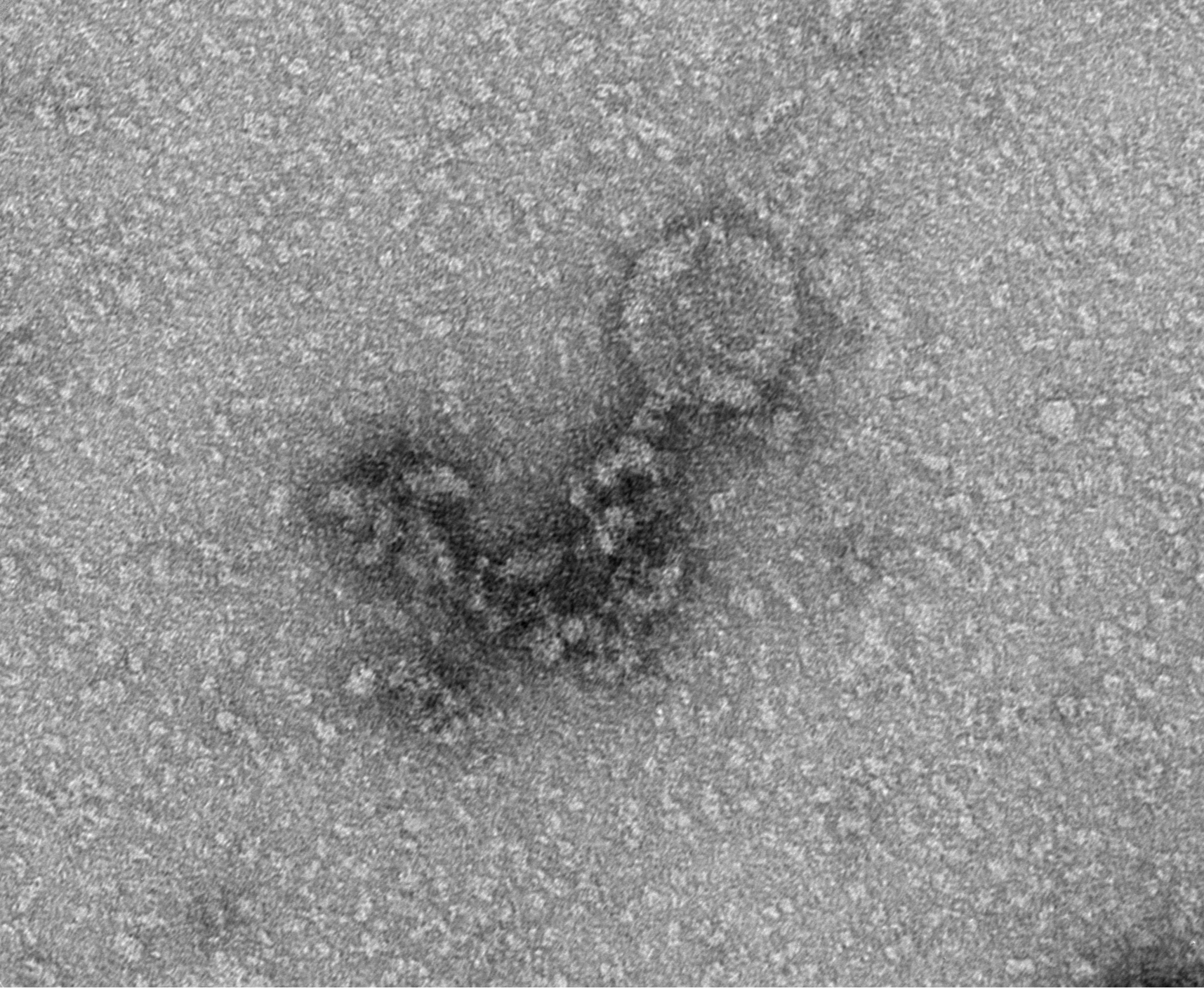


X-54_TEM1-PHG1_S1_009.tif
X-54_Ravel
TEM1-PHG1_S1
Cal: 0.145603 nm/pix
9:38:52 a 05/30/24
TEM Mode: Imaging
Microscopist: MLD

HV=80.0kV
Direct Mag: 67000x
UMB EMCIF

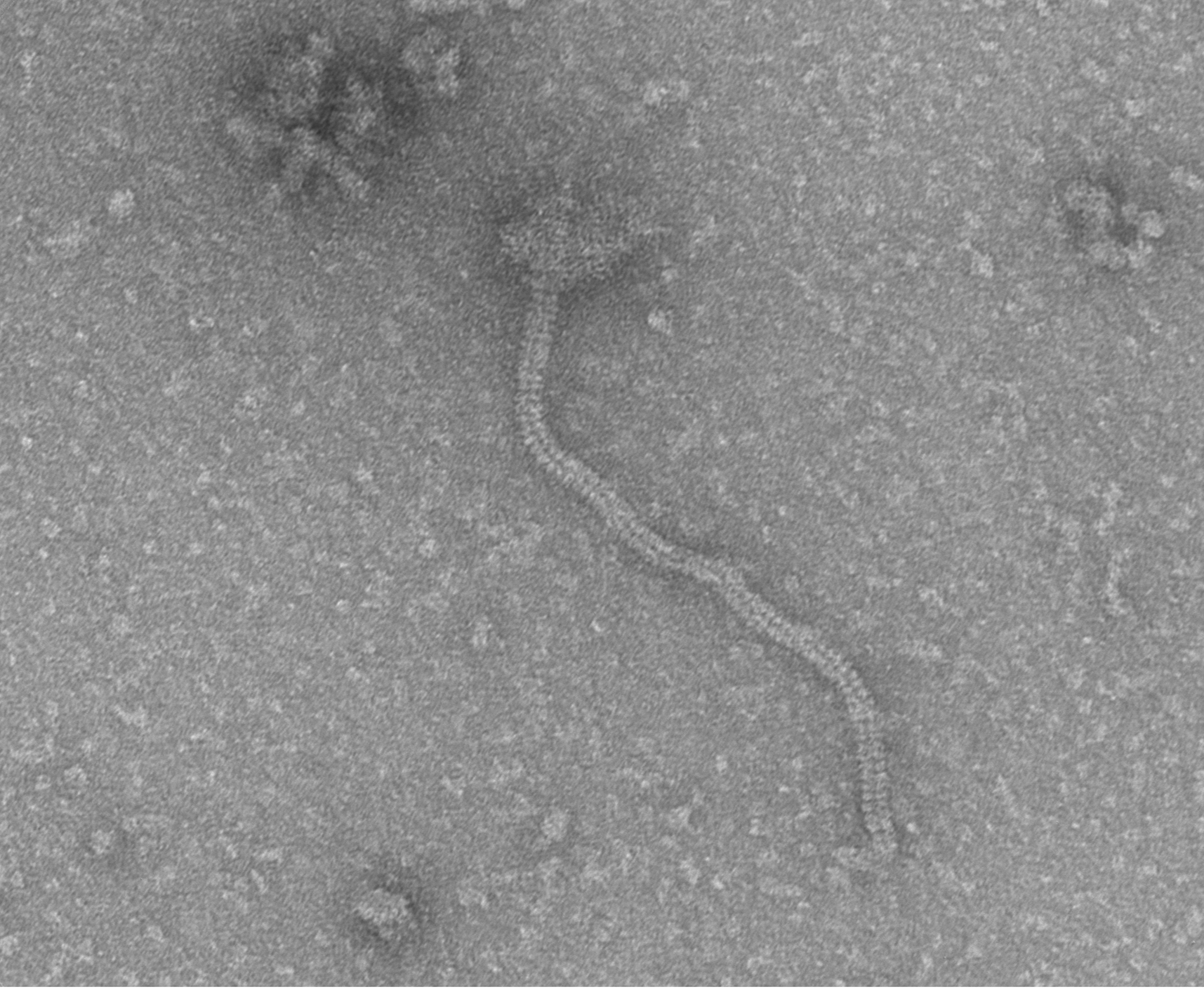

X-54_TEM1-PHG1_S1_032.tif
X-54_Ravel
TEM1-PHG1_S1
Cal: 0.145603 nm/pix
10:27:25 a 05/30/24
TEM Mode: Imaging
Microscopist: MLD

20 nm
HV=80.0kV
Direct Mag: 67000x
UMB EMCIF
